# Deploying photons for communication within neuronal networks

**DOI:** 10.1101/2021.08.02.454613

**Authors:** Montserrat Porta-de-la-Riva, Adriana Carolina Gonzalez, Neus Sanfeliu-Cerdán, Shadi Karimi, Sara González-Bolívar, Luis Felipe Morales-Curiel, Cedric Hurth, Michael Krieg

## Abstract

Deficiencies in neurotransmission lead to neurological disorders or misinterpretation of perceived threats. To restore defects in cellular communication, we developed a synthetic, photon-assisted synaptic transmission (PhAST) system. PhAST is based on luciferases and channelrhodopsins that enable the transmission of a neuronal state across space, using photons as neurotransmitters. We demonstrate the ability to overcome synaptic barriers and rescue the behavioral deficit of a genetically engineered glutamate mutant with conditional, Ca^2+^-triggered photon emission between two cognate neurons of the *Caenorhabditis elegans* nociceptive avoidance circuit. We also deploy these ingredients for asynaptic transmission between two unrelated cells in a sexually dimorphic neuronal network. Functional PhAST could sensitize otherwise poorly responsive males to touch and hence expand the behavioral repertoire. Our study, thus, establishes a powerful framework for complex photon-based communication between neurons in a living animal, that can readily be expanded to synthetic neuronal networks, organoids or non-invasive brain-machine interfaces.

---

A major overarching challenge in applied neuroscience is to establish control over spatiotem-poral signalling within the brain. Optogenetics (*1*) is a promising strategy to control neuronal activity by exploiting orthogonal light-activated ion channels (*2*) that are ectopically expressed in target neurons. However, in vertebrates—including humans—the light needs to be delivered via skull-implanted light sources, which emit potentially harmful intensities (*3*) in order to reach target neurons in deeper brain layers. Due to scattering of light in dense brain tissue, a light source must be close to target neurons in order to achieve cell- or even circuit-specific activity (*3*). Recently, bioluminescence-driven optogenetic effectors were introduced for blue-light informed, trans-cellular signal trans-duction (*4*), a strategy that bears extraordinary potential for establishing prosthetic neurotransmitters in living animals (*5*). However, the implementation of photons as transcellular signals remained challenging, primarily because of the low quantum yield inherent to bioluminescence and the resulting difficulty in recruiting sufficient numbers of active channels for postsynaptic depolarization in a cell-specific manner.

Here, we capitalized on the simple genetics and known structural and functional connectome of the model organism *Caenorhabditis elegans* to establish a genetically encoded, cell- and thus circuit-selective optogenetic sniper strategy to control neuronal activity at the synaptic level without the need for external light delivery. In order to achieve photon-amplified synaptic transmission (PhAST), we targeted the expression of calcium-dependent synthetic luciferases (enhanced Nanolanterns (*6*), eNLs) as conditional quantum emitters to presynaptic neurons and combined them with postsynaptic localized high-photocurrent channelrhodopsin mutants (*7*). Our ultimate goal was to complement a chemical synaptic transmission defect engineered into the well-characterized nociceptive avoidance circuit of *C. elegans* (*8*).

We took advantage of a neuronal model circuit defective in glutamatergic neurotransmission resulting from the lack of *eat-4*, a vesicular glutamate transporter responsible for packaging these neurotransmitters in synaptic vesicles (*8–10*). ASH is a polymodal nociceptor that responds to mechanical nose touch and makes direct synaptic connections with AIB and AVA neurons in an *eat-4*-dependent manner (*8, 11, 12*) (Fig. S1A,B). We compared the behavioural response to external nose touch delivered by an eyebrow hair to the tip of the nose of the animal (where sensory endings of mechanical nociceptors are located; Fig. 1A) in wildtype and mutant animals (Fig. S1). In agreement with previous work (*9*), we found that wildtype animals responded to 70% of nose contacts (Video 1, Fig. S1B), while *eat-4(ky5)* mutant animals only responded to 2% of touches (Fig. S1C,D).

**Figure 1.**
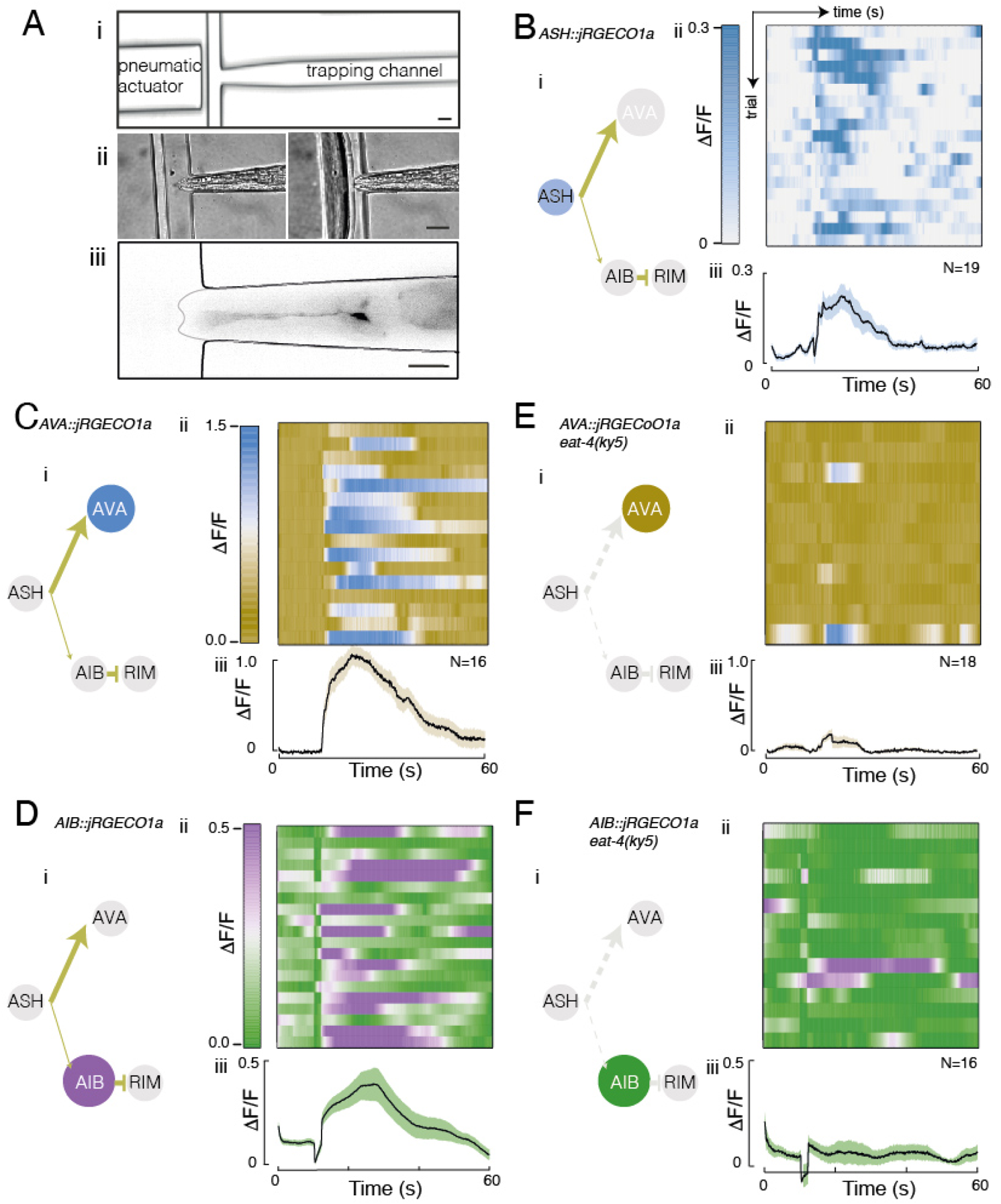
A nose-touch defect derived from mutations in synaptic transmission. A (i) Layout of the central part of the microfluidic Trap’N’Slap with (ii) representative pictures of a trapped animal before and during mechanical stimulation. (iii) Representative image of an animal expressing the calcium indicator GCaMP in ASH. Scale bars = 20 µm. B-F i) Schematic of the ASH nociceptive avoidance circuit with nodes indicating presynaptic (ASH) and postsynaptic (AVA, AIB, RIM) neurons; edges are colour-coded according to neurotransmitter (yellow, glutamate). Grey dashed edges correspond to disrupted, thus inactive neurotransmission in the eat-4 mutant. Node coloured according to the kymograph lookup table. Edge thickness reflects the number of connections between neurons. ii) Normalized and baseline-subtracted kymograph of neuronal cell body intensity versus experimental time. A 2-s stimulus was delivered after 10 s. Each row (N) is derived from a different stimulation. iii) Average, normalized fluorescence intensity of the Ca2+ indicator in (B) ASH, (C,E) AVA, and (D,F) AIB in (B-D) control and (E,F) glutamatergic mutant animals (eat-4(ky5)). Mean ± standard deviation is shown. See Methods for details.

In order to specifically observe the activation of ASH and the transmission of neuronal signal from ASH to AVA and AIB (Fig. 1), we designed a microfluidic device that delivers mechanical stresses to the nose while simultaneously measuring calcium activity in ASH and interneurons (Fig. 1A, Fig. S2). We called this device Trap’N’Slap. The Trap’N’Slap contains a pneumatic actuator (*13*) that drives a deformable polydimethylsiloxane diaphragm (Fig. 1a, Fig. S2C, Video into an immobilized animal, permitting high-resolution fluorescence imaging. We characterized this deformation as a function of pressure and confirmed that our method visibly deforms the nose of a trapped animal (Fig. 1A, Fig. S2D,E and Video 2). Next, we loaded animals expressing the genetically encoded fluorescent calcium indicators GCaMP (*14*) or jRGECO1a (*15*) in the ASH sensory neuron into the Trap’N’Slap; both dyes reproducibly underwent an increase in signal intensity upon pneumomechanical nose touch (Fig. 1B, Fig. S3A).

Having shown that ASH is specifically activated in our micromechanical device, we next engineered jRGECO1a specifically into AVA and AIB interneurons (Fig. 1C,D) using promoters previously described (*8, 16, 17*) (Methods), with the goal of following signal transmission from the sensory to the interneuron layer. After a pneumatic nose touch delivered for 2 s, both AVA and AIB robustly activated with Ca^2+^ response dynamics that greatly exceeded the stimulus duration (Fig. 1C,D). When we presented the same stimulus to the glutamate-deficient *eat-4(ky5)* animals, we still observed ASH responses (Fig. S3B), but AVA and AIB failed to respond with discernible Ca^2+^ dynamics (Fig. 1E,F), indicating that chemical communication between ASH and AVA/AIB on the synaptic level was effectively broken. Together, our pneumatically actuated microfluidic devices and calcium-imaging system constituted a framework for our subsequent efforts to optically restore and follow the flow of information in a neuronal circuit through PhAST.

Next, we expressed light-sensitive ion channels as effectors in the postsynaptic interneurons AIB and AVA (Fig. 2a). We generated transgenic animals expressing channelrhodopsin2-**H**a**RDC**ore (ChR2-HRDC) with the previously characterized H134R (*18*) and D156C (*19*) mutations, yielding a light-gated channel with an unprecedented photon-current relationship and improved surface expression (*7*). We selectively expressed ChR2-HRDC in AVA using an intersectional genetic strategy (*16*) and in AIB with the *npr-9* promoter as previously (*8, 17*) (Fig. 2B,C; Methods). We confirmed the functionality of the channelrhodopsin by recording the escape response after illuminating individual animals transgenic for ChR2-HRDC in both AIB and AVA (Fig. 2D, Fig. S4, Video 6, 7) with and without the all-trans retinal (ATR) photosensitizer (Fig. 2E). We carefully titrated decreasing light levels and extrapolated the data with a binary logistic regression model (Methods) to estimate the response probability at the lowest light intensities (Fig. 2F). With this approach, we inferred that animals still responded at intensities below 1 fW/*µ*m^2^. For comparison, because a single photon carries an energy of 1e-19 J, responses at the lowest light intensities used here were evoked with fewer than 10.000 photons*·*s*^−^*^1^*µ*m*^−^*^1^. Importantly, no activity was recorded in AVA neurons that did not express ChR2-HRDC (data not shown) and in AVA neurons that were not supplemented with ATR (Fig. 2E, Suppl. Text), consistent with light triggering ChR2-HRDC activity and concomitant neuronal depolarization. We also recorded the escape response in animals lacking ASH specific or systemic glutamatergic signalling; AIB response was dependent on *eat-4* in downstream neurons, whereas AVA response was not (Fig. S4). In summary, we established the most sensitive neuronal system for light-driven behavioural responses (Fig. 2F) in *C. elegans* reported to date.

**Figure 2.**
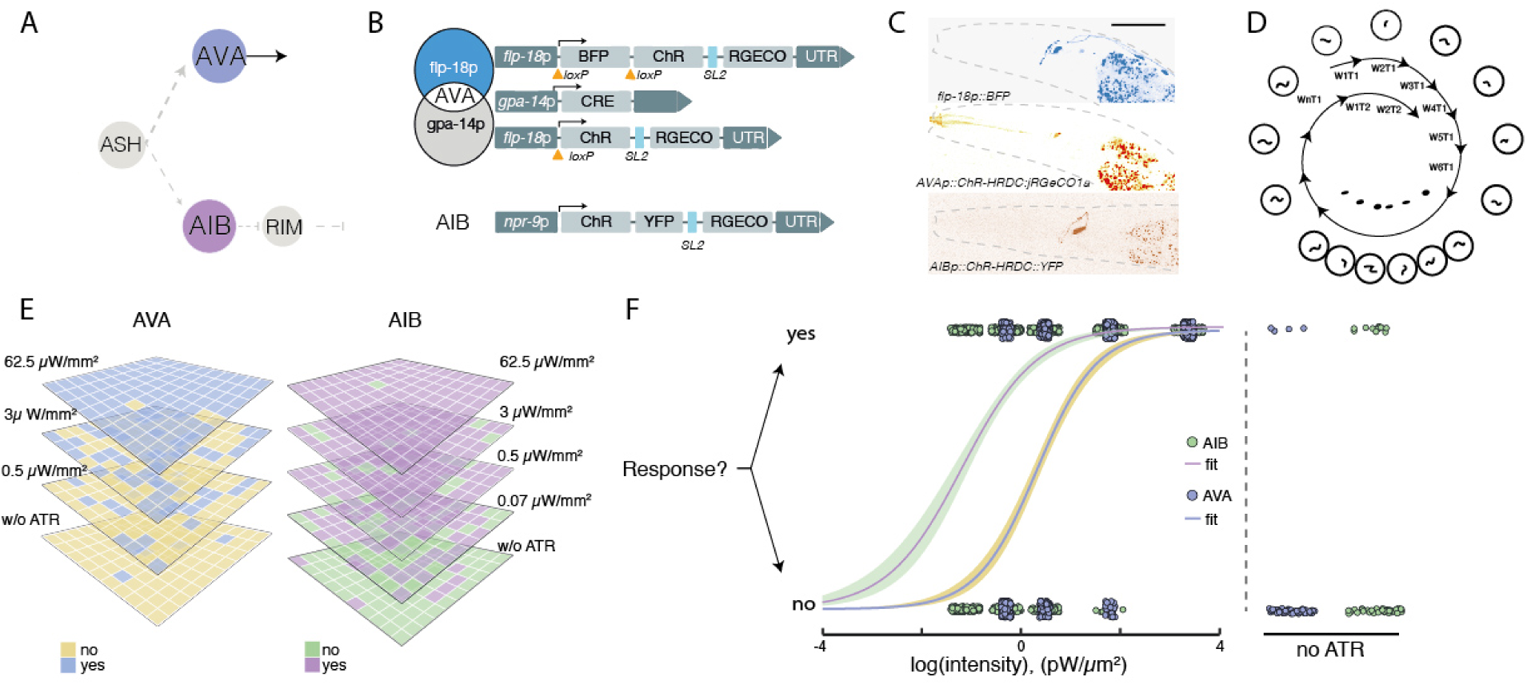
Postsynaptic light sensitivity of a new combination of ChR2 mutants. **A** Schematic of the ASH nociceptive avoidance circuit with ultra-sensitive light-gated ChR2s in AVA and AIB. Grey dashed edges correspond to disrupted neurotransmission in the *eat-4* mutant. **B** Genetic strategy for cell-specific targeting of ChR2-HRDC to AVA and AIB. AVA was targeted using an intersectional strategy employing promoters *flp-18* and *gpa-14*, which exclusively overlap in AVA. Successful recombination removes the loxP-flanked BFP and brings ChR-HRDC::jRGECO1a under *flp-18p* control. ChR-HRDC expression in AIB was achieved with the single *npr-9* promotor as described (*17*). **C** Representative confocal microscopy pictures of AVA expressing BFP and the red-shifted Ca^2+^ indicator jRGECO1a before (upper) and after (middle) CRE-mediated recombination. The lower image depicts AIB expressing ChR2-HRDC::YFP. Scale bar = 30 µm. **D** Experimental routine for interrogating light-sensitive behaviour. A single worm (Wx) per plate was stimulated with blue light once (T1) before trialing the next plate with a different animal. Ten rounds of one stimulation constitute a single dataset. In total, 30 animals were tested, each 10 times. **E** Representative outcome of the behavioural response to blue light at the indicated intensities of animals expressing ChR-HRDC in AVA or AIB in the absence or presence of the photosensitizer ATR. **F** Behavioural avoidance response curve as a function of light intensity. Solid line is a binary logistic regression of the no/yes response. For control experiments, ATR was omitted from the food source and individual animals were illuminated at the maximum light intensity of 2.4 mW/mm^2^. N=300 stimulations of 30 animals per condition.

To obtain a genetically encoded light source that functionally interacts with light-gated ion channels, we engineered a conditionally light-emitting luciferase into ASH mechanosensory neurons as a source of quantum emitters. eNLs (*6*) are chimeras that carry a luciferase moiety (luc) fused to a fluorescent protein that selects the colour of the emitted photons. To facilitate a good spectral match with downstream channelrhodopsin while maximizing energy transfer from luciferase to the fluorescent protein, we chose mTurquoise2 (*20*) as the photon emitter (Fig. 3A).

**Figure 3.**
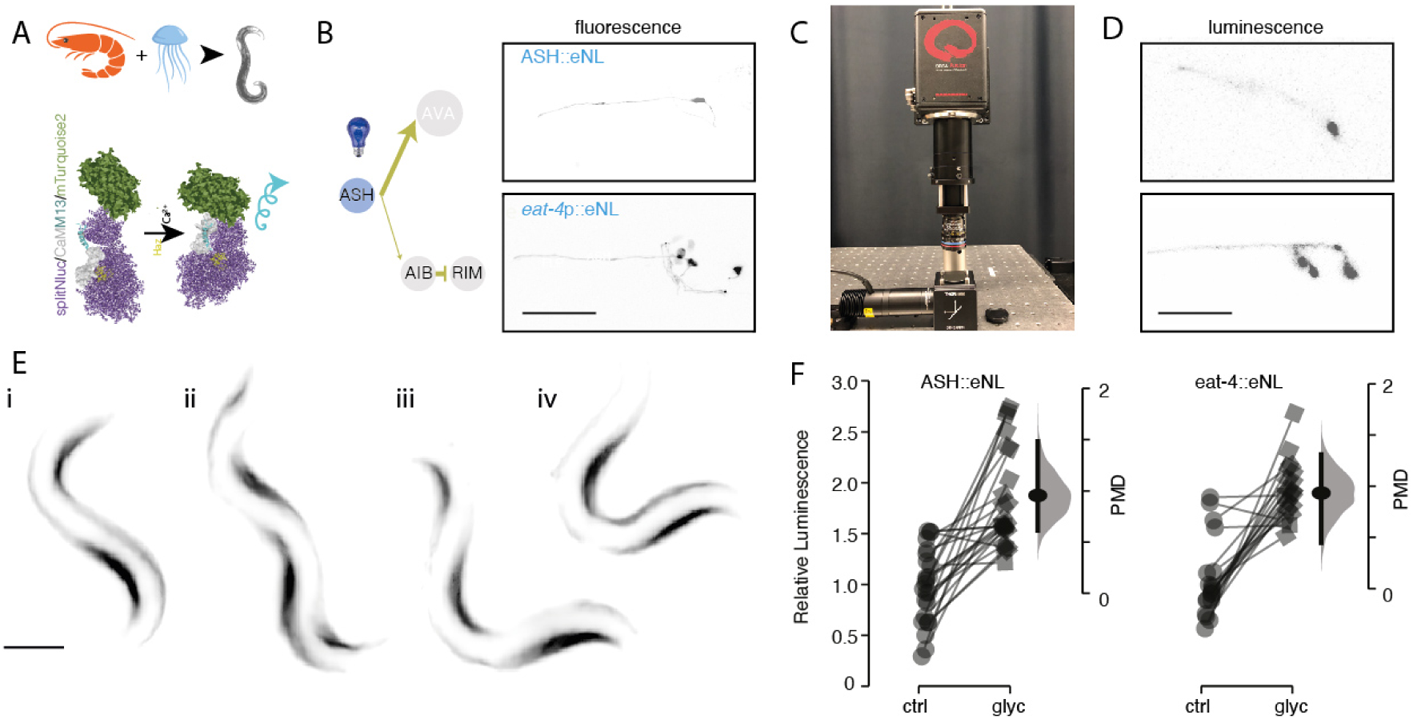
A calcium-triggered synaptic photon emitter. **A** Schematic of the working principle of the switchable luciferase. A luciferase fused to mTurquoise2 (eNL (*6*)) is reconstituted upon Ca^2+^ binding and oxidizes a cofactor (yellow) in order to emit light. **B** Schematic of the circuit for eNL expression in ASH with representative commercial spinning disk confocal microscopy of ASH (top) and all glutamatergic neurons (bottom) in *C. elegans*. Scale bar = 50 *µ*m. **C** LOwLIght microscope with an optimized optical axis and low-noise photodetectors. **D** Bioluminescence emitted by ASH (top) and all glutamatergic neurons (bottom) acquired on the custom LOwLIght microscope. Exposure time = 1 s. Scale bar = 50 *µ*m. **E** Luminescence micrographs of a crawling sequence recorded with an animal expressing calcium-sensitive eNLs in body-wall muscles, representative for 8 out of 10 videos. Images recorded with 1 s of exposure time on the LOwLIght microscope. See also Video 8. Scale bar = 100 *µ*m. **F** Neuronal activation by 0.1 mM glycerol triggers photon emission through Ca^2+^ entry. Measurements are paired with and without glycerol. The floating axis to the right indicates a bootstrapped distribution of the paired mean difference (PMD), with the horizontal bars indicating the 95% confidence interval. P-values were *p <* 1*e −* 12 for both comparisons as derived from a two-sided permutation t-test.

Importantly, the luciferase in eNL is split by a calcium-sensing domain to achieve conditional photon emission in the presence of the high Ca^2+^ concentrations that are typical for neuronal activation (*21*). Given the estimated resting Ca^2+^ concentration of 60-90 nM in ASH (*22*), we chose a calcium sensor domain with a K*_d_* of 250 nM in order to maximize the sensitivity and dynamic range of our eNL (*6*).

We first expressed a codon-optimized eNL under the ASH-specific *sra-6* promoter (*23*) (Fig. 1A) as well as the *eat-4* promoter, which is active in all glutamatergic neurons, including ASH nociceptors (Fig. 3B). We confirmed strong cyan fluorescence in the heads of animals under external blue light excitation (Fig. 3B). However, due to the limited photon budget of luciferases in the absence of high intracellular calcium (*6*), it was impossible to visualize light emission resulting from luciferase’s intrinsic activity using standard optical microscopes (data not shown). To overcome this limitation, we built an improved LOw-LIght microscope with an optimized optical axis, high-power objectives, and a single photon-sensitive camera chip (Fig. 3C). In this configuration and with novel chemical cofactors delivering higher quantum yield (hikarazine (*24*)) and bio-availability (fluorofurimazine (*25*)), we visualized photon emission from both of our ASH and glutamate transgenes, demonstrating that luciferases can emit light *in situ* (Fig. 3D). We also visualized photon production by eNLs upon an increase in calcium influx in body-wall muscles and observed light emission by the eNLs on the contracted body-wall muscles of *C. elegans* during unconstrained animal locomotion (Video 8; Fig. 3E). These data establish that our eNLs increase their quantum yield and emit photons more efficiently in the presence of calcium.

To further visualize how ASH activity and the concomitant increase in intracellular calcium (Fig. S3) induces a quantum emission, we performed a calcium-imaging experiment in the Trap’N’Slap under mechanical stimulation in our LOw-LIght microscope (Fig. 1A, Fig.3). However, even with the technical improvements in microscopy and cofactor chemistry described above, the obtained signal was very faint, especially at short exposure times (Fig. 3D), due to the low ‘off’ activity of the Ca^2+^ dye. This limitation precluded functional imaging with the eNL under mechanical stimulation. To achieve our goal of observing an increase in the quantum yield of photons triggered by neuronal depolarization, we resorted to a luminescence plate reader capable of recording and counting relative luminescence levels (Methods). We first recorded baseline luminescence in ASH-specific and broadly expressed eNLs in glutamatergic neurons. Then, we added 0.1 mM glycerol, which repels *C. elegans* (*26*), and measured luminescence 3 s later. ASH is the main— and so far only—polymodal neuron known to evoke a cellular response and calcium increase upon glycerol-mediated osmolarity changes (*26, 27*). Strikingly, we detected a significant and similar increase in photon yield for two strains expressing an eNL exclusively in ASH or, more broadly, in glutamatergic neurons (*p <* 1 *·* 10*^−^*^8^, *N >* 20, permutation t-test (*28*); Fig. 3F). Taken together, these experiments demonstrate that calcium-induced photon emission under physiological conditions is fast and reproducible in freely behaving animals.

Having shown that photon emission can be triggered by stimulation of neuronal activity in presynaptic compartments, we expressed the eNL and ChR2-HRDC together in the same worm, supplemented the animal’s diet with both co-factors (ATR and Hikarazine), and assayed the prosthetic circuit’s efficiency in complementing the genetic *eat-4(ky5)* disruption of the glutamatergic signalling pathway from ASH→AVA/AIB (Fig. S5A). We first immobilized individual animals in the trapping channel of Trap’N’Slap and delivered a pneumatic stimulus to the nose. In agreement with our previous experiment (Fig. S1), animals carrying all transgenes but lacking the two cofactors did not exhibit a calcium increase in response to mechanical nose touch (Fig. S5B). However, when these animals were fed both cofactors, we detected a robust increase in AVA and, to a lesser extent, AIB activity after nose touch (Fig. S5C,D). This result motivated us to ask whether the observed signal transmission from the sensory to the interneuron layer is sufficient to drive reversals in the nose touch avoidance behaviour. We thus counted the number of times that an individual animal with glutamatergic deficits displayed an escape response upon nose touch with an eyebrow hair, as an indicator for a functional reconstitution of the nociceptive avoidance circuit in presence of the required cofactors. Even though we registered more behavioural responses on the population level for *eat-4* mutants that were supplemented with Hikarazine and ATR than untreated mutant controls, the average log odds ratio of detecting a positive response versus no response in eah individual *eat-4* mutant did not depend on cofactor presence (Fig. S5F). Thus, we were not able to detect an effect of our treatment on the single-animal level. We conclude that despite detecting an increase in Ca^2+^ activity, we failed to observe rescue of nose touch avoidance behaviour responses (Fig. S5E,F).

Because many neurons downstream of ASH are glutamatergic (e.g. AIB and RIM (*29*)), we reasoned that a lack of systemic glutamate signalling interferes with successful reconstitution of the nociceptive avoidance response. We thus established a conditional CRE/lox strategy to obtain a cell-specific knockout of *eat-4* restricted to ASH sensory neurons. We first flanked exons 1 and 2 with two loxP sites using CRISPR/Cas9 (Fig. S6A,B) and confirmed that neither Ca^2+^ signalling in AIB (Fig. S6C) nor avoidance behaviour (Fig. S6E,H) were significantly affected by the genomic loxP sites or by the expression of CRE by itself (Fig. S6F,H). We then coexpressed CRE and confirmed successful recombination with a fluorescent reporter of CRE activity (*30, 31*) (Fig. S6B,D). Successful excision of *eat-4* by a conditional CRE recombinase is expected to delete the two described transcription start sites and to lead to loss-of-function of glutamate signalling through ASH and a nose touch phenotype. As expected, we consistently observed a loss of nose touch avoidance behaviour (Fig. S6G,H) consistent with a defect in ASH signal transmission. In agreement with previous results (*32*), of the various mechanosensors that activate upon mechanical nose touch, ASH contributes to more than 60% of the total responses recorded to nose touch. Thus, a conditional allele can be used to interfere with synaptic transmission in ASH to downstream interneurons.

We then engineered a cell-specific *eat-4* knockout mutant with the ASH-restricted eNL (Fig. 2 and AVA::ChR2-HRDC (Fig. 2E) and visualized Ca^2+^ signals in AVA after stimulation in the microfluidic chip (Fig. 4A). Without the critical cofactor for the eNL, no calcium dynamics corresponded with the mechanical stimulus from the pneumomechanical device in AVA or AIB (Fig. 4B,C). In contrast, a robust Ca^2+^ increase in AVA related to the pneumatic stimulus to the nose occurred after incubation with a high concentration of Hikarazine (Fig. 4D,E). As in worms harbouring the systemic glutamate defect (Fig. S5), in these ASH(*eat-4*) knockout animals, AIBdid only respond occasionally to the imposed stimulus (Fig. 4F).

**Figure 4.**
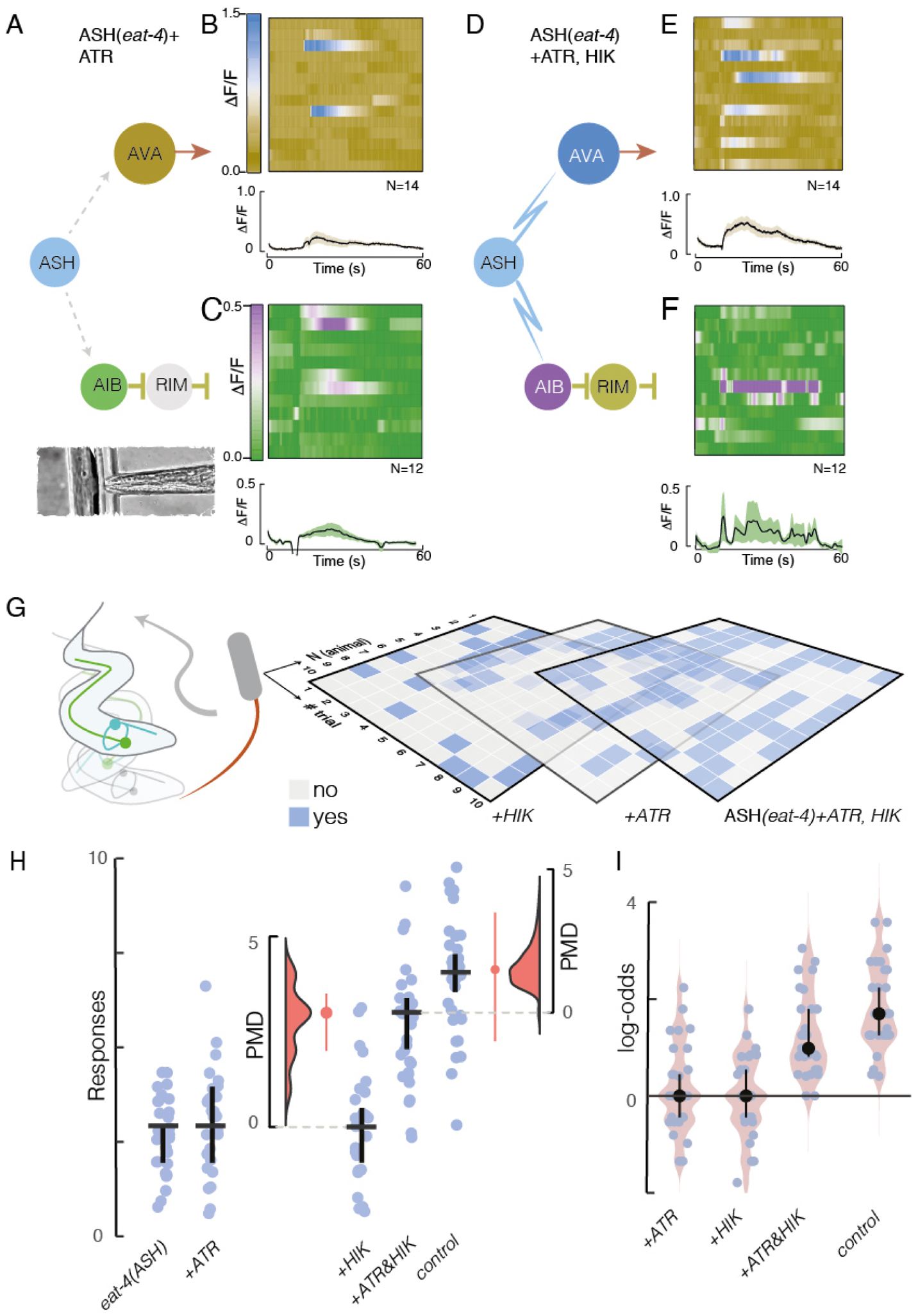
PhAST rescues nociceptive avoidance behavior and postsynaptic neuronal activity. **A-F** Calcium recordings from the indicated neurons of ASH-conditional *eat-4* mutant worms trapped inside the microfluidic chip (A-C) without and (D-F) with prior exposure to the cofactors hikarazine (HIK) and ATR. (A,D) Schematic of the circuit with nodes colour-coded according to the look-up table of the kymographs and edges coloured according to the neurotransmitter used (yellow, glutamate; red, acteylcholine; grey, mutant condition; blue, photons). (B,C,E,F) Stacked kymographs of individual Ca^2+^ recordings from (B,E) AVA and (C,F) AIB in the conditional ASH mutant (ASH(*eat-4*)) with (E,F) and without (B,C) cofactors. Plots below the kymographs depict mean*±*standard deviation (N=number of recordings). **G** Schematic of the behavioural experiment with three representative grid plots of the response of ASH-conditional *eat-4* mutant animals in the presence and absence of cofactors. **H** Rescue experiments in the conditional *eat-4* background. A vertical jitter was applied for display purposes to differentiate individual datapoints. Horizontal bar indicates the median, vertical bar indicates 95% confidence interval of the median (N=30 animals). Floating axes indicate the bootstrapped distribution of the paired median difference and its 95% confidence interval. **I** Log-odds ratio of detecting a positive response in the indicated animals compared to untreated mutant animals. Median*±*95% confidence interval. Control = wildtype animal without a defect in glutamatergic signalling.

We next sought to determine whether PhAST from the sensory to the interneuron layer rescues the ASH-specific defect and elicits a nociceptive avoidance response in our behavioural paradigm (Fig. 4G). In the absence of both cofactors, baseline behavioral activity was similar to that in the ASH(*eat-4*) mutant (Fig. 4G). We then carried out the nose-touch assay 10 times on 30 animals harbouring three alleles of the same transgenes (1000 touches per condition; Table S1, Fig. 4, Fig. S7). In addition, we tested an eNL that was specifically targeted to the synapses in ASH through a *sng-1* fusion (*33*) (Fig. S8A,B) and detected a consistently higher response probability in animals supplemented with ATR and hikarazine (Fig. S8C,D). With the most efficient transgene, the behavioural response was close to that of worms without *eat-4* defects (Fig. 4I). Statistical modeling of the response rate (Methods) suggested that the odds of rescuing avoidance in mutants is up to 20 times higher with PhAST than without it in average(Fig. 4I). Taken together, these experiments establish that photons can be used to encode and transmit the activity state between two neurons within a neuronal circuit.

We next sought to test a gain-of-function experiment and asked if PhAST can sensitive a behavioral response by wiring two neurons that normally do not form synaptic connections. The connectome of *C. elegans* suggests various sex-related differences between hermaphrodite and male individuals (*34*). In particular, no synapses have been found between ASH and AVA in males (Fig. 5A,B) (*34*), suggesting a sexually dimorphic behavioral nociceptive avoidance response. In-deed, males do not avoid nose touch with an eyebrow hair as efficiently as hermaphrodite animals (Fig. 5C) and AVA in *C. elegans* males did not respond with an increase in Ca^2+^ to mechanical stimulus delivered through the Trap’N’Slap device (not shown). We thus sought to wire the connection between ASH and AVA with the aim to sensitize and ‘feminize’ the response of males to mechanical nose touch. To achieve this, we performed our PhAST experiments in males in absense and the presence of the necessary cofactors ATR and hikarazine. Strikingly, in presence of both cofactors, *C. elegans* males responded almost indistinguishable to hermaphrodites (Fig. 5C) and showed up to 10 times higher odds ratio than the average of untreated males. Together, this shows that PhAST is able to sensitize a sexually dimorphic circuit for nociceptive avoidance behavior and thus functionally wire an unconnected pair of neurons.

**Figure 5.**
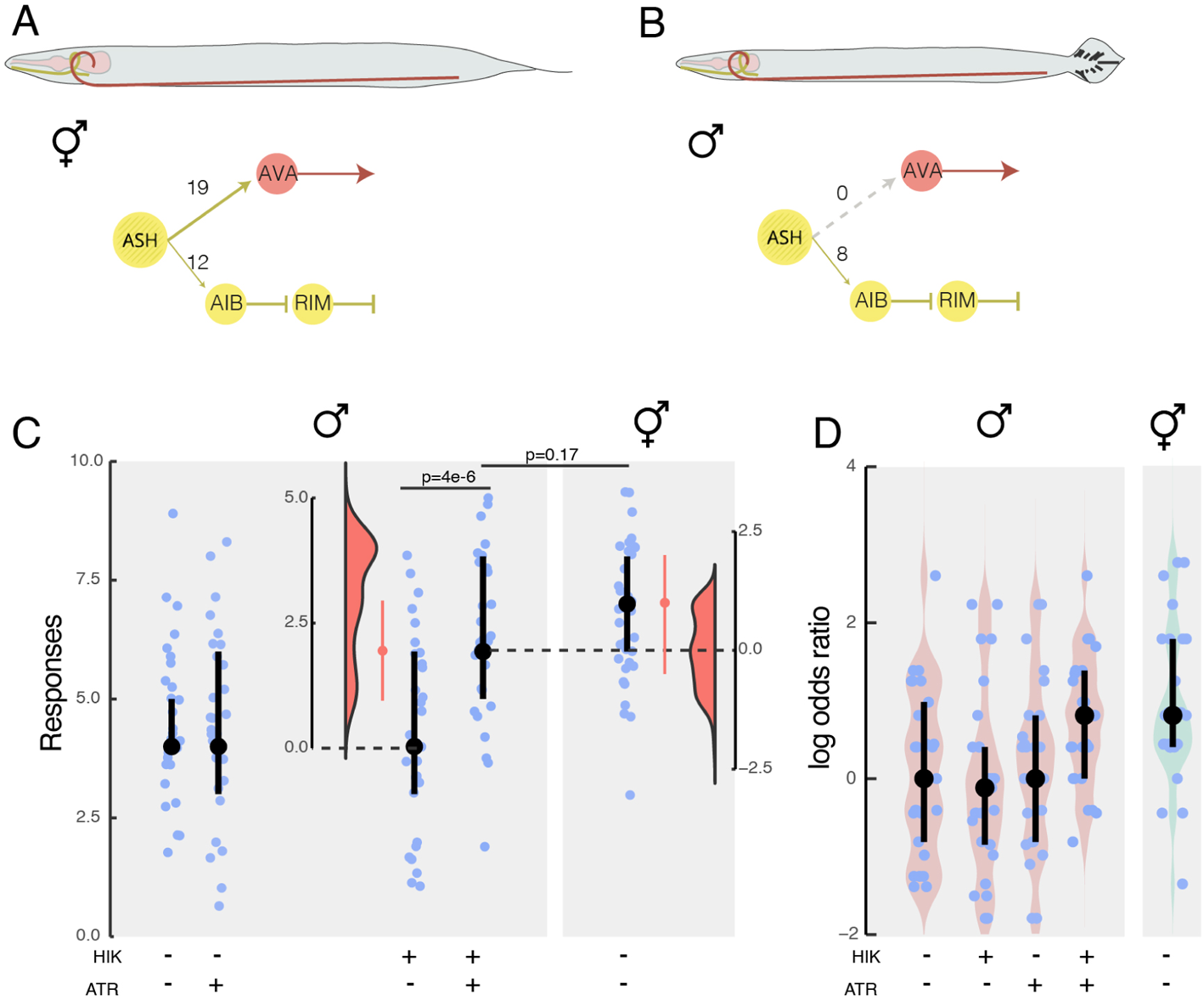
Feminizing a sexually dimorphic behavior in males with PhAST. **A, B** Sketch of a (A) hermaphrodite and a (B) male with its corresponding nociceptive avoidance circuit. Numbers on the edges indicate the sex-specific numbers of synapses detected in electron micrographs (*34*). **C** Individual responses of male animals expressing the PhAST system in presence and absence of the ATR and hikarazine cofactors, compared to untreated hermaphrodites of the same genotype. A vertical jitter was applied for display purposes to differentiate individual datapoints. Dot indicates the median, vertical bar indicates 95% confidence interval of the median (N=30 animals). Floating axes indicate the bootstrapped distribution of the paired median difference and its 95% confidence interval. p-values derived from Wald test statistics above indicated combinations. **D** Log-odds ratio of detecting a positive response in an individual animal of the indicated condition compared to an untreated random male animal. Individual males were compared to randomized individuals. Median*±*95% confidence interval.

In summary, here we replaced the mechanism for endogenous chemical neurotransmission between a sensory neuron and a pair of interneurons with a genetically encoded, photon-assisted synaptic transmitter (PhAST) system and showcased its ability to overcome a genetically constructed synaptic barrier. Then we use PhAST to wire anatomically unconnected neurons with a behavioral consequence. These experiments constitute the first demonstration that light can be genetically encoded as the transmitter of a state variable between two cells. Our research therefore complements luminopsin, a self-illuminating microbial rhodopsin that is a fusion between channelrhodopsin and a luciferase, which was previously employed as an inhibitory construct to suppress the activity of coelenterazine-exposed transgenic neurons (*35*).

Will this approach be universally applicable across the neuronal network? Based on optimistic estimates of photon production by eNLs and the activity states of ChR2-HRDC (SI Text), we assume that not more than a dozen channels are open at the same time in a single neuron. Given the extraordinarily high input resistance of *C. elegans* neurons (*36*), the simultaneous opening of a few ion channels likely depolarizes the neuron by tens of millivolts (*37*), which is sufficient for signal propagation in isopotential neurons (*38*). The next challenge will be to couple unrelated circuits and to generate or suppress synaesthetic interactions (*39*) between sensory neurons and downstream interneurons or to construct synthetic, self-actuating networks based on artificial neuromuscular communication (*40*). Future improvements in bioluminescent enzymes, light-gated ion channels, and subcellualar targeting will enable unprecedented optical control over neuronal function, non-invasively and with extraordinary specificity and precision.

## Acknowledgments

We thank the NMSB and SLN labs for discussions and suggestion throughout the work and for use of their microscopes. We thank the ICFO BIL and NFL for support with animal maintenance and SU8 lithography, respectively. We thank Cori Bargmann, Shawn Xu, Sander van der Heuvel, Alexander Gottschalk, Peter Askjaer, Thomas Kirkland and the CGC (National Institutes of Health - Office of Research Infrastructure Programs (P40 OD010440)) for providing reagents; Yves Janin, Pau Gorostiza, Robert Kittel, and Anna Kim for discussions and suggestions on bioluminescence, optogenetics and microfabrication; and Pau Gorostiza and Peter Askjaer for critical comments on the manuscript.

## Funding

MK acknowledges financial support from the ERC (MechanoSystems, 715243), HFSP (CDA00023/2018), Spanish Ministry of Economy and Competitiveness through the Plan Nacional (PGC2018-097882-A-I00), FEDER (EQC2018-005048-P), “Severo Ochoa” program for Centres of Excellence in R&D (CEX2019-000910-S; RYC-2016-21062), from Fundació Privada Cellex, Fundació Mir-Puig, and from Generalitat de Catalunya through the CERCA and Research progam (2017 SGR 1012), in addition to funding through H2020 Marie Skłodowska-Curie Actions (754510 to AG) and Mineco (FPIPRE2019-088840 to NS).

### Author contributions

MPR: animal husbandry, molecular biology, CRISPR, CRE recombination, optogenetic and nociceptive experiments, data analysis, and manuscript writing. ACG: microfluidics, calcium imaging, bioluminescence imaging, and the first manuscript draft. NS: behavioural assays, molecular biology. CH, SG, SK: microfabrication, design, simulations. LFCM: bioluminescence imaging, microscopy. MK: Study concept, acquisition of funding, data analysis, software programming, and manuscript writing.

### Code and Material availability

All reagents produced are freely available upon request to the corresponding author. All data needed to evaluate the conclusions in the paper are present in the paper and/or the Supplementary Materials. Some strains will be deposited to the CGC. Scripts developed supporting the analysis can be accessed under Gitlab::NMSB.

## Supplementary Data: Functional photon communication within neuronal networks

### 1. Supplementary Text 1: Estimates for photon budget

A recent estimate in the number of available channelrhodopsin necessary for neuronal depolarization of mammalian neurons was previously estimated to be in the order of 1 million molecules (*1*). How do our result compare with this estimate? We calculated the anticipated and required photon budget to depolarize the postsynaptic neuron sufficiently to evoke signal propagation via light-activated channels. We first estimated how many photons we will obtain given a certain luciferase concentration and synapse volume. Using published values for quantum yield (10/s) (*6*) and typical overexpression concentrations (10*^−^*^6^M or 3000 molecules) in a synapse with a radius of 0.5*µ*m (*≈*0.5fL), we expect to obtain approx. 30000 photons/s/synapse. Since a typical calcium transient in ASH in our experiments lasts about *>>*5s (Fig. 1) and ASH forms 27 synapses with AVA and AIB (Fig. S1, Ref. (*11*)), we thus expect to have *≈*4.5*10^6^ photons available for stimulation, or 2pW. Because the synaptic cleft is less then the wavelength of the light, we assume there is neither absorption nor scattering, such that this value corresponds to a photon flux of 0.7µW/mm^2^ through the postsynaptic membrane halfspace. Because not all photons are emitted in direction of the target cell, this value is overestimated by a factor that depends on the area overlap in the synaptic contact.

To calculate the relative photon absorption *E* of the ChR2 in the membrane we apply Lambert-Beer’s law, which relates the path length *d* (thickness of the membrane/channelrhodopsin), the concentration of absorbers *c* and extinction coefficient *E* (1-photon absorbance cross section, 50000 L mol*^−^*^1^ cm*^−^*^1^ for ChR2; (*4*)). With reasonable values of the surface area of a synapse of 1*µm*^2^ and an estimated density of 190 molecules/*µ* m (*19*), we estimate the concentration of ChR2-HRDC of 62*µ*M within the postsynaptic membrane. Subsequently, according to Lambert-Beer (*E* = *c∗E∗d*) the relative photon absorption is E=3.6e-6, meaning that the intensity of the light after passing through the membrane with ChR2 is *I*_0_ *∗* 10*^E^* and, consequently, *≈*4 out of 1 million photons will be absorbed. Because the photon absorption vs. activation ratio of ChR is 0.7 (*4*), we estimate the quantum efficiency of the system is 2.5 *·* 10*^−^*^4^ %. Taken together, given the 4.5*10^6^ photons emitted from the Nanolantern, we expect to activate *>*20 channels during a typical stimulation. Because ChR2-XXL is 200 times more sensitive than ChR2 (*19*), the real number is likely much higher.

How many open channels are necessary to depolarize the neuronal membrane for a given amount in a given time. We consider published values of the resting potential of −50mV for AIB (*8*) and −30mV for AVA (*7*) and asked how many charges are necessary to depolarize AVA for 20mV, a hypothetical value to activate voltage gated Ca channels. Note, AIB is not known to express low threshold T-type calcium channels (e.g. cca-1) that would activate at lower membrane potentials (−30mV for NMJ, (*8*)), whereas AVA expresses both, T-type and L-type voltage gated Ca channels (*9*). Given an input resistance of 5GOhm, a current of 5pA is necessary to do so.

Assuming a specific capacitance of 1*µF/cm*^2^ and a synaptic radius of 500nm, a minimal amount of *≈* 4000 sodium ions would be required to raise the potential of about 20mV. Due to the overlap of sodium entry and potassium exit (during action potential), we consider that 4 times more ions are required (*10*). With a published conductance of 750fS for ChR2-XXL (*19*), a single channel would conduct 300000 ions/s at such an electrochemical driving force, taking about 100ms to depolarize the synaptic compartment sufficiently. Because we are using an ultrasensitive ChR, called ChR2-HRDC, with an improved conductance and membrane stability, in principle we would only need one active channel to produce a depolarization of 20mV to elicit secondary responses critical for signal propagation.

### 2. Supplementary Text 2: CRE recombinase expression

In a first attempt to overlap expression of *eat-4* and the CRE recombinase, we used the *octr-1p* promoter, which was described to be expressed in a restricted number of cells in the head (*11*). After confirming a restricted expression and overlap with ASH (Fig. S9A) of the *octr-1* promoter cloned we tested the response to nose touch of worms coexpressing the floxed *eat-4* allele and *octr-1p*::CRE. Animals failed to respond consistently to nose touch and frequently lost the mTagBFP2 marker in somatic tissue, displaying ubiquitous recombination pattern in all cells, which we attributed to the reported expression in spermatheca and/or germ line.

To avoid the spurious recombination in the germline, we performed a split-CRE (*12*) approach in which we targeted the 3.8kB *sra-6* promoter (*23*) together with *gpa-13* promoter and the split form of CRE (Fig. S9B). We observed successful reconstitution of the CRE activity in 2-4 cells in 90% of the animals, which correlated with a decrease in nose touch response in animals coexpressing the floxed *eat-4* allele (Fig. S9B). This confirmed that *eat-4* was effectively excised with the split CRE, without affecting other tissues.

We reasoned that splitting the CRE enzyme might result in a recombination efficiency *<*100%. In order to increase the recombination efficiency in ASH, we surveyed various promoters that exclusively drive expression in glutamatergic neurons involved in nose touch. Since we also use a CRE/lox strategy to obtain AVA-specific ChR2 expression, we looked for promoters with an overlap on ASH and AVA. Recently, Schmitt et al (*16*) proposed an intersectional strategy using a *gpa-14p*::CRE and a *flp-18p* transgene that generates an AVA restricted transgene expression after recombination. Among other neurons in the head where *gpa-14* is expressed (Fig. S9C), ASH is the only neuron involved in nose touch. We confirmed recombination in ASH with coexpression of an *sra-6p*::GCaMP transgene, that leads to overlap in red and green channel. We then assayed nose touch and found a significant decrease in the reversal rate, similar to the other transgenes tested and what was observed before for ASH ( (*32*), Fig. S9C). We attributed the residual nose touch response to other nociceptive avoidance neurons, e.g. FLP. Indeed, when we deleted *eat-4* in FLP using a *des-2p*::CRE construct, in addition to ASH, we found a strong reduction in the nose touch response (data not shown).

### 3. Supplementary Text 3: Choice of Luciferase and Channel-rhodopsin

We reasoned that the spectral overlap between the luciferase emission and ChR2 absorption is critical for the function of the system and thus we only considered a combination of eNL with blue-activated ChR2. However, using red-shifted ChR2 like Chrimson or ChRmine, we anticipate that luciferases like Antares are superior. Even though the common firefly luciferase emits photon that peak at Chrimson absorption, we were unable to observe a large photon production in transgenic animals for firelfy luc.

We first considered the general ChR2-H134R as photosensitizer, but discarded it due to the low photon current (not shown). We then turned our attention to the high photocurrent ChR2 bearing the mutation C128S;L132C;H134R (*7*) (hereafter termed ChR2-triple) and generated transgenic animals expressing ChR2-triple in AVA. In young adult animals, we observed a strong response at 2.4mW/mm^2^, but a fast habituation to repetitive stimuli. Moreover, older (day 2 onwards), animals lost their ability to respond to blue light, due to neuronal degeneration, visible as loss of AVA. We suspected that a continuous depolarization in presence of ATR led to this effect. We thus generated the double mutant ChR2-HRDC, which we employed in this study for downstream experiments due to its unprecedented ability to repetitively drive behavior in *C elegans* at extreme low light intensities.

### 4. Methods

#### *C. elegans* culture

Nematodes were cultivated on NGM plates seeded with *E. coli* OP50 bacteria using standard protocols (*17*). Unless otherwise stated, age synchronized young adult hermaphrodites, except for experiments related to Fig. 5, were used for the experiments.

### Molecular biology

Unless otherwise specified, all plasmids used for this study were assembled using the Gibson assembly method. Sequences are listed in table S4.

#### 4.1 Expression of ChR and jRGECO1a//ChR::YFP and jRGECO1a in AIB

Sequence for *npr-9* promoter was obtained from (*17*). pNMSB34 was constructed by directed mutagenesis on the universal MosSCI plasmid pNMSB29, which contained *npr-9p*::ChRTriple::SL2::jRGECO1::*unc-54* 3’UTR with primers specified in Table S5. pNMSB37 was constructed inserting YFP from pSX-317 (gift from Shawn Xu, (*8*)) between ChR2-HRDC and SL2::jRGECO1a by Gibson assembly.

#### 4.2 Expression of ChR and jRGECO1a in AVA

An intersectional strategy ensured cell-specific expression in AVA. We first introduced a loxP-mTagBFP2-stop-loxP::ChR-SL2-jRGECO1a construct using the universal MosSCI method (*19*). The universal MosSCI plasmid pNMSB6a was built as follows: mTagBFP2::*tbb-2* 3’UTR, ChR Triple (Bergs et al, 2018) and SL2::jRGECO1a fragments synthesized by TWIST BIOSCIENCES were assembled using a 4-fragment Gibson assembly into pNMSB6 which already contained *flp-18p* and *unc-54* 3’UTR. To facilitate conversion, we co-expressed *gpa-14p*::CRE, effectively turning AVA from blue to red, indicative for a succesful jRGECO1a expression.

The ChR triple turned out to be toxic for AVA in adult animals (see supplementary text 3). To convert ChR triple to ChR2-HRDC, a CRISPR was performed with crRNAs and a homology-directed repair template as indicated in Table S5. All reagents were purchased from IDT and conditions for injection were: 12.5 *µ*M each crRNA, 2 *µ*M crRNA for *dpy-10*, 27 *µ*M tracrRNA, 6 *µ*M Cas9, 0.5 nM *dpy-10* ssODN and 1.75 nM homology repair template.

#### 4.3 AVA::CRE

For *gpa-14p*::CRE, pNP259 plasmid described in (*16*) was used. To confirm cell-specificity of the recombination, we established the *gpa-14p*::CRE in the SV2049 strain (gift from Sander v d Heuvel), in which successful recombination can be followed by a BFP-mCherry switch of fluorescent proteins (*30, 31*). We consistently found recombination in AVA and also in ASH.

#### 4.4 Expression of calcium sensitive, enhanced Nanolantern (eNL) in ASH, glutamatergic neurons and muscles

eNL250 was synthesized by TWIST BIOSCIENCES using a *C. elegans* codon optimized version of mTurquoise2 and the Ca^2+^ 250 eNL described in (*6*) and cloned into pMINI-T (Invitrogen). pN-MSB16 (*sra-6p*::eNL250::*unc-54* 3’UTR) was built replacing the ORF in pNMSB14 (2.0 kb *sra-6p*::RCaMP1h::*unc-54* 3’) by NL250 from pMINI-T-eNL250. pNMSB17 (*eat-4p*::eNL250::*unc-54* 3’UTR) was constructed amplifying the *eat-4* promoter (table S4) from genomic DNA, and assembling it in a 3-fragment Gibson assembly reaction with the pMINIT-NL250 and a backbone containing *unc-54* 3’UTR. pNMSB40 was bluit by replacing the mCherry driven by the *myo-3* promoter in pCFJ104 by eNL250.

#### 4.5 Targeting eNL to pre-synaptic regions in ASH

To build pNMSB26 (*sra-6p*::*sng-1*::NL250::*unc-54* 3’UTR) the sequence for *sng-1* was synthesized by TWIST BIOSCIENCES and cloned into pMINI-T. A 2-fragment Gibson assembly was performed to introduce the ORF for *sng-1* between the *sra-6* promoter and NL250 in pNMSB16 with a flexible linker between them.

#### 4.6 Split CRE expression in ASH

For cell-specific expression of CRE exclusively in ASH, we followed an intersectional strategy that involves a split CRE (*12*) construct under the control of two promoters that are exclusively co-expressed in ASH. The N-terminal fragment of the split CRE (aa 1-244) construct was synthesized by TWIST BIOSCIENCES flanked by NaeI and EagI restriction sites, and subsequenetly cloned into a vector containing a *gpa-13* promoter and *unc-54* 3’UTR (pNMSB5). The synthetic C-terminal fragment of the split CRE (aa 245-345) was cloned into a 3.8 kb *sra-6* promoter (*23*) containing the *unc-54* 3’ UTR (pNMSB3) by directed mutagenesis with primers in table S5.

#### 4.7 *octr-1p*::CRE

*octr-1p* was amplified from genomic DNA (Table S4) and cloned in pNMSB7 together with the three intron CRE::tbb-2 3’UTR in pDD282 (*22*). The fragment *octr-1p*::CRE::tbb-2 3’UTR was then moved to the universal MosSCI vector pNMSB28.

#### 4.8 ASH:jRGECO1a

pNMSB72 was constructed by replacing in pNMSB6a the regions corresponding to *flp-18* promoter, mTagBFP2 and ChR by the 2.0 kb *sra-6* promoter and the miRFP670 ORF (addgene-plasmid-79987, see table S4).

### Transgenesis

Transgenic animals were generated by microinjection of varying amounts of DNA according to standard protocols and the conditions (amount/composition) indicated in table S3. For extrachromosomal arrays, plasmid DNA was mixed with DNA ladder to a maximum DNA load of the mix of 100 ng/*µ*L. The integration of the extrachromosomal array was performed using UV/TMP method. In brief, late L4 or young adult animals carrying the array were picked onto a NGM plate without OP50. These animals were fed TMP at a final concentration of 50 *µ*g/mL for 30 minutes. Then, they were UV irradiated for 30 seconds at 4.5mW*·*cm^-2^ (250nm peak wavelength; *≈*130mJ) and expanded for 3-4weeks before selection. Three independent integrated lines were recovered whenever possible.

#### 4.9 Generation of the floxed *eat-4*

Floxed *eat-4* allele was generated by CRISPR following the protocol described in (*23*). In a first editing step, a loxP site was introduced before the first aminoacid using two different crRNAs and a HR-template as indicated in Table S5. In a second step, the loxP site was introduced after the second exon by means of the crRNAs and HR templates indicated in Table S4. All reagents were purchased from IDT. Injection conditions were: 12.5 *µ*M each crRNA for *eat-4*, 2 *µ*M crRNA for *dpy-10*, 27 *µ*M tracrRNA, 6 *µ*M Cas9, 0.5 nM *dpy-10* ssODN and 1.75 nM eat-4 homology repair template.

#### 4.10 CRE transgenes

The recombination efficiency was determined using a CRE reporter strain (*30, 31*) (gift from Sander v d Heuvel), carrying a transgene with floxed blue fluorescent protein (BFP), followed by mCherry. Successful recombination results in a blue to red color switch in cells with active CRE enzyme. To follow recombination, candidate animals were immobilized on agar pads and imaged on a DragonFly Spinning Disk confocal microscope with 405nm laser excitation (BFP) and 594 nm laser excitation for mCherry.

### Microfluidic chip device design and operation: Trap’N’Slap

All designs were made in AutoCad 2019 and UV printed with a maskless aligner (MLA, Heidelberg Instruments). The loading chamber and trapping channel geometry was copied directly from Ref (*13*). The height of the channel was 50*µ*m to accommodate day one adults. Three different actuators were designed with 50*µ*m height, 50, 100 and 200*µ*m width and 15 *µ*m membrane thickness. The wider actuators can be deflected further at the expense of response time. In the ultimate design, the squeezing actuator was set at a width of 200*µ*m.

#### 4.11 Analytical calculation of diaphragm deflection

To characterize actuator deflection as a function of back pressure, we connected the pressure inlet to the OB1 8bar pressure channel (ElveFlow) and increased the back pressure in 50kPa steps while taking images of the inflated diaphragm. Diaphragm deflection was measured in ImageJ.

As shown in eq. 1, maximum deflection of an elastic rectangular membrane with module of elasticity of (*E*), is linearly correlated with pressure *P* acting on its surface with thickness of (*t*), width of (*w*) and height of (*h*). The coefficient of *α* empirically depends on edge condition and mechanical properties of the material (*25*).

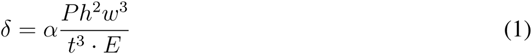

The Young’s modulus of a Polydimethylsiloxane (PDMS (1:10), (Sylgard 184 by Dow Corning)) membrane with 15 microns thickness is set 1.6 MPa based on studies (*13, 26*) on thickness-dependent mechanical properties of PDMS membranes. Since applying lower pressure and achieving larger deflection in our chip is ideal, we used 1:15 PDMS mixing ratio. Lower amount of reagent means less molecular binding, which leads to more flexible membrane and higher deflection and the results are shown in Figure S2. The Young’s modulus for this mixing ratio is 1.4 MPa. The width and height of the membrane are 200 and 50 microns, respectively. The coefficient of *α* is set 0.0034 for our actuator’s edge condition which is only fixed from the part that is bonded to a cover slip (*25*).

#### 4.12 Finite Element Analysis (FEA)

In order to have a prediction that is more accurate we performed numerical simulation based on FEA. In real case, since the hydrostatic pressure applied in the channel also deforms the sidewalls of the actuator, therefore we considered the surrounding walls in the simulation. The actuator is simulated with Ansys workbench (2021 R1). The material is set as a PDMS block as obtained from a 1:15 mixing ratio cured at 85°C for two hours with the specification of Young’s modulus of 1.4 MPa and 0.5 Poisson’s ratio with tensile strength 2.24 MPa (*26*). The model is meshed using structured hexahedral grids (Fig. S2) to reduce the mesh size and thus computational cost while maintaining the appropriate grid quality. We conduct mesh independence studies in CFD (computational fluid dynamics) to make sure that the results we get are due to the boundary conditions and physics used, not the mesh resolution. Mesh independency is assessed based on total deformation. Average cell size is sequentially reduced until the displacement difference is independent from the grid size. The obtained results are independent of the mesh size above 127320 nodes (not shown). Thus, the cell size corresponding to the case with 127320 nodes is chosen for the numerical investigation. The actuator was studied under three pressure rates (0, 150, and 350 kPa). Both results from the Eq. 1 and simulation showed that lower thickness results in steeper slope, which requires lower pressure to apply in the channel to achieve the desire deflection. To optimize the dimensions of the actuator, different width, thickness and height were studied. Since the channel geometry is set by the dimensions of the animals and cannot be changed, thickness and elasticity of the diaphragm are the major design variables permitted. Lower thickness plays an important role to increase the deflection but lower than 10 microns is challenging from fabrication point of view. Measurements confirmed that the length of the channel had a negligible effect on membrane deflection.

#### 4.13 Device Fabrication

The fabrication of the molds was undertaken in-house as a single layer process using standard SU8 soft-lithography techniques (*27*). The 4-inch wafers were cleaned with Piranha cleaning standard process to remove trace amount of organic contamination and residuals. In brief, we first applied a 5um thick adhesion layer to reduce lift-off of the patterned structure during device fabrication and cured it under direct UV exposure for half an hour. Then SU8-50 was coated on the substrate and baked at 65 and 95 degrees. The design which is created with AutoCAD is converted to the format of MLA software (CleWin). The pattern is printed on the substrate and post baked before developing in SU8 developer for 6 minutes and rinsed with propanol. The mold is ready after hard baking 2 hours on 135 degrees. After fabrication, molds were vapor-phase silanized in chlorotrimethylsilane to prevent adhesion of the PDMS to the substrate. A 15:1 mixture of Sylgard 184 prepolymer/curing agent was degassed (*≈*30min in vacuum desicator) and poured onto the silanized molds. After settling, the PDMS/wafer were baked at 85°C for two hours. Devices were then cut using a scalpel, lifted off and punched with a biopsy punch (0.75mm). Coverslips (Menzel Gläser #1.5) were cleaned in a 2-Propanol bath for 10 min and properly rinsed in ddH_2_O. PDMS/glass bonding was performed with a 15s plasma activated treatment (Plasma Asher PVA TePla 300) followed by an annealing bonding process of 10 min at 120°C in a hotplate. Quality of the seal was tested manually *in-situ* with a syringe connected to the actuator inlet.

#### 4.14 Animal loading and experimental setup

The procedure of animal insertion into the trapping channel has been described in detail elsewhere (*28*). In brief, to load individual in the chip, three to four synchronized day one adult worms (*17*) were picked from an NGM plate containing OP50 bacteria and transferred to a NGM plate without food to rid themselves from bacteria. Then, these animals were placed in a 15 *µ*L droplet of physiology buffer (145 mM of NaCl, 2 mM of Ca_2_Cl, 1 mM of MgCl_2_, 5 mM of KCl, 10 mM Hepes and 25 mM of Glucose, with a pH 7.4). Using a stereo dissecting scope at 60x total magnification (Nikon SMZ25), the animals were aspirated into a SC22/15 gauge metal tube (Phymep) connected to a 3 mL syringe (Henke Sass Wolf) with a PE tube (McMaster-Carr) prefilled with physiology buffer. The loading tube was inserted in the inlet port of the device, while a gentle pressure onto the plunger of the syringe released the animals into the loading chamber. The pillars in the loading chamber act as a sieve and slow down the animals, such that they can be oriented head-first for efficient mechanical stimulation. The animal was positioned such that 20µm of the nose protruded into the flush channel, ready to accept a mechanical stimulus. If the worm was entering the channel in the wrong direction, the syringe was pulled gently at the same time another syringe connected to the outlet tube was used to apply back pressure to help orient the worm. For the experiments where the furimazine analog corresponding to the hydrolysis of Hikarazine-108 (*24*) was used, a similar procedure was performed with the exception that the animal was preincubated for 10s in a 2 µl droplet of the cofactor, followed by aspiration in the PE tube filled with physiology buffer.

#### 4.15 Mechanical stimulation and calcium imaging

The animal loaded device was then positioned on a Leica DMi8 and the stimulation channel was connected to a piezo-driven pressure controller (OB1-MK3, Elveflow). With a 10x/0.3 objective lense the animal was positioned within the field of view and then with a 40x/1.1 water immersion lens in place to focus on sensory neuron ASH or interneurons AVA and AIB. The neurons were identified based on the specific expression of jRGECO1a and anatomical location at the posterior pharyngeal bulb. Image acquisition was performed with a Lumencor SpectraX light engine (cyan LED with 470 nm bandpass cleanup for GCaMP and green/yellow LED through a 555 nm bandpass filter for jRGECO1a; 5% transmission corresponding to *≈*0.5mW at the sample plane) and a filtercube containing 570 nm edge dichroic mirror and 515/15nm emission filter and 595/20 nm emitter for GCaMP and jRGECO1a, respectively. Videos were captured with 10 Hz with a Hamamatsu Orca Flash 4.3 for 10-60 seconds using HCImage. A masterpulse was used to synchronize the camera acquisition with the light exposure. Exposure time was set to 85ms. The camera SMA trigger out was used to synchronize the stimulation protocol setup in ElveFlow sequencer software prior to the imaging routine. The sequence consisted of 100 prestimulus frames, two seconds of a pressure step of 300kPa and 48s poststimulus acquisition.

#### 4.16 Image analysis

Images were preprocessed in ImageJ and then imported into python to extract signal intensity using in-house procedures. In short, the image sequence was cropped to a small area surrounding the cell body of the neuron of interest and a smooth filter was applied. The resulting image stack was analysed with a python script and the signal intensity was extracted based on the calculation of the 10th percentile to measure the background and the 90th percentile to measure the neuron intensity. After background subtraction, signal intensity was normalized to the first 100 frames (before the mechanical stimulus was applied).

### Behavioral assays

#### 4.17 Nose touch assays

Plates for nose touch assays were prepared as follows: 10 mL of NGM were poured to 5.5 cm plates and allow to dry over night at room temperature. The next day, 100 *µ*L of an overnight OP50 culture diluted 1:1 in LB were spread onto the plates allowed to dry and grown over night at room temperature. Plates were either directly used or stored at 4°C until needed. Worms for the assay were transferred as L4 and assayed the next day as young adults. For the assays, a special picker was used with an eyebrow hair at the tip. The eyebrow hair was placed in front of the worm so that it could freely contact the hair (*30, 32*). A positive event was counted when, upon the contact of the tip of the nose with the hair, the worm reacted moving backward.

#### 4.18 Animal preparation for optogenetics

Animals were cultivated in the dark at 20°C on NGM with OP50 bacteria with or without all-trans retinal (ATR) (*18, 32*). Plates containing ATR were prepared by spreading 100 *µ*L of OP50 culture mixed with ATR (0.1 mM final concentration) onto 3.5 cm plates containing 3.5 mL of NGM. About 16 h before the experiments, L4 worms grown on regular NGM plates were transferred to fresh assay plates. For measurements, worms were illuminated with blue light (467-499 nm) at the specified light intensity, under a 2x objective on a Nikon SMZ25 stereomicroscope. Duration of illumination was defined manually and lasted for 1 s. Every single worm was illuminated 10 times with a minimum interstimulus interval of 1 minute. Any observable backward locomotion during or directly after (1 s) a blue light pulse was scored as a response. The incident power of the excitation light was measured with a microscope slide powermeter head (Thorlabs, S170C) attached to PM101A power meter console (Thorlabs).

#### 4.19 Animal preparation for rescue experiments

About 20-24h before the experiments, L4 worms grown on regular NGM plates were washed off the plates with S-medium complete and finally transferred to 2mL eppendorf tubes containing 250*µ*L of S-medium complete (*33*) supplemented with 0.05% Triton and 1% DMSO. 10*µ*L of OP50 five times concentrated was added as food supply (with or without ATR). Concentrated solution of the luciferin corresponding to Hikarazine-108 (its O-acetylated form, provided by Yves Janin, Institute Pasteur) was obtained by performing its hydrolysis using a mixture of DMSO and ethanol containing 37% hydrochloric acid (10 eq.) as previously described (*24, 41*). Where indicated, Hikarazine (*24*) was added as the substrate for bioluminescence at a final concentration of 0.4 mg/mL. Final ATR concentration was 0.1 mM.

Worms were incubated at 20°C for 20-24h in the dark and constant rotation. After that time, the liquid was transferred to plates in the same ATR conditions and worms transferred to fresh plates once the liquid was dry. After 2h recovery on the plates, nose touch test was performed in the dark with a 590nm bandpass filter (Thorlabs) to block the blue component of the white light used in the stereomicroscope.

#### 4.20 Statistics of the behavioral assay

Behavioral data is scored as a binary yes/no (1, 0) response as a result of a mechanical stimulation. Thus, the obtained data is binomially distributed with a single categorical independent design variable representing the predictor (treated vs non-treated; luminescent vs dark; wild type vs mutant) of the response. We modeled the outcome of each experiment with a general linear model after binary logistic regression such that

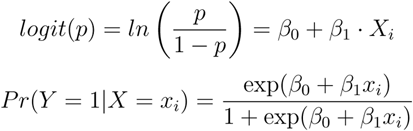

or which simply says that the probability of getting a yes response for the categorical variable *x_i_* = 1 (e.g. wild type animals) is an odds ratio. In the above case, *β*_0_ is the value of the transformed outcome variable when the predictor is equal to zero (mutant, non-treated, dark), *β*_1_*x* describes the increase in odds of finding a positive response for *x_i_* = 1. For a continuous independent variable (e.g. light intensity), the odds increase by *exp*(*β*_1_) for each unit increase in *x*_1_. We plot the log-odds ratio of obtaining a positive response to a mechanical stimulus in the rescued conditions in Fig. 4 and Fig. S5 with respect to the mutant condition. Significance of the parameters was tested using the Wald chi-squared statistics according to *z*^2^ = (*β*^^^*_j_/*SE(*β*^^^*_k_*))^2^.

Similarly, the optogenetic data is a binomially distributed with a single continuous predictor variable, e.g. light intensity. To extrapolate the light response at low intensities beyond the accessible experimental parameters, we fitted a generalized linear model to the raw counts of individual responses.

Wherever indicated, we resorted to p-value independent statistical comparison by estimating the paired median difference (PMD) between two sample distributions and concluded that a large effect existed if the 95% confidence interval of the median PMD does not overlap with zero. This is indicated as a floating axis in selected comparisons of the data (Fig. 3e, 4H, S1, S5, S6, S7, S8). The PMD distribution was calculated by bootstrapping a sample containing 40 datapoints for at least 100 times. For each bootstrapped sample the difference in µ1 and µ2 was calculated.

### Bioluminescence imaging

Bioluminescence imaging was impossible on a standard compound microscope and we redesigned a compressed optical path to enhance quantum efficiency in extreme lowlight conditions (LoLi). The details are described elsewhere, but in short, a 100mm tube lens (Applied Scientific Imaging) is used to focus light collected by a 40x/1.25 silicon immersion objective (Olympus) directly onto a Hamamatsu Orca-Fusion camera (C14440-20UP) with no additional optical elements in place. Exposure time was adjusted to maximize acquisition frame rate in expense of signal/noise ratio and generally kept below 1s. Images were denoised using convolutional neural networks (*34*) trained on experimental data (publication in preparation). During training, by iteratively minimizing the loss function through stochastic gradient descent, the network weights were optimized to improve the image reconstruction. The dataset consisted of paired ground truth images and noisy images, and was augmented via random change, spin, and rotation to improve the inference quality and avoid overfitting.

To image animals expressing the eNL confined to the body wall muscles, worms were placed onto a 1% agarose pad prepared in a glass slide. Here, worms are able to perform body bend but do not crawl. Animals were then shortly incubated with 20 mM of the FFz cofactor in physiology buffer (*25*) or Hikarazine-108 (*41*) and covered by placing a coverslip. Imaging of bending worms was performed for *≈*60s by using a 1s exposure time.

#### 4.21 Quantification of luminescence in Microplate reader

The relative light units of ASH eNL and *eat-4* eNL strains were measured with a microplate reader via glycerol-mediated chemical stimulation. An average of 100 animals per strain were placed in triplicates in a white flat bottom 96-well plate. Three endpoint measurements were consecutively made per well: baseline luminescence without cofactor addition, basal photon emission after cofactor addition (0.1 mM Hikarazine) and neuronal photon emission after addition of 0.1 mM glycerol. Values were subtracted to the baseline and a ratio after and before worms were supplemented with glycerol was calculated. Values are represented as Mean *±* SD.

### Confocal microscopy

Fluorescence images were taken using an inverted research microscope (Nikon Eclipse Ti2) equipped with a spinning disk confocal microscope (Andor DragonFly 502, Oxford Instruments) on top of an active isolation table (Newport). A 60x/1.2 NA CFI Plan Apo VC water immersion objection and Andor Sona camera were used. mTagBFP2 was excited using the 405 nm laser, 30% power intensity and transmitted through a 445/46 nm emission filter. Exposure time varied between 30-100 ms, depending on the strain to image. mTurquoise2 was excited using the 445 nm laser, 30% power intensity and transmitted through a 478/37 nm emission filter. Exposure time varied between 100-200 ms depending on the strain imaged. YFP was excited using the 514 nm laser, 20% power intensity and transmitted through a 552/41 nm emission filter. Exposure time was 30 ms. jRGECO1a was excited with 488 and 561 nm lasers, 30% power intensity each and transmitted through a 594/43 nm emission filter. Exposure time varied between 100-200 ms depending on the strain imaged. mCherry was excited with a 561 nm laser, 10% power intensity and transmitted through a 647/63 nm emission filter. Exposure time was 30-100 ms. GCaMP/GCaMP7 were ex-cited with a 488 nm laser, 40-80% power intensity respectively and transmitted through a 521/38 nm emission filter. Exposure time was 30-200 ms respectively.

## 5 Supplementary Figures

**Figure S1.**
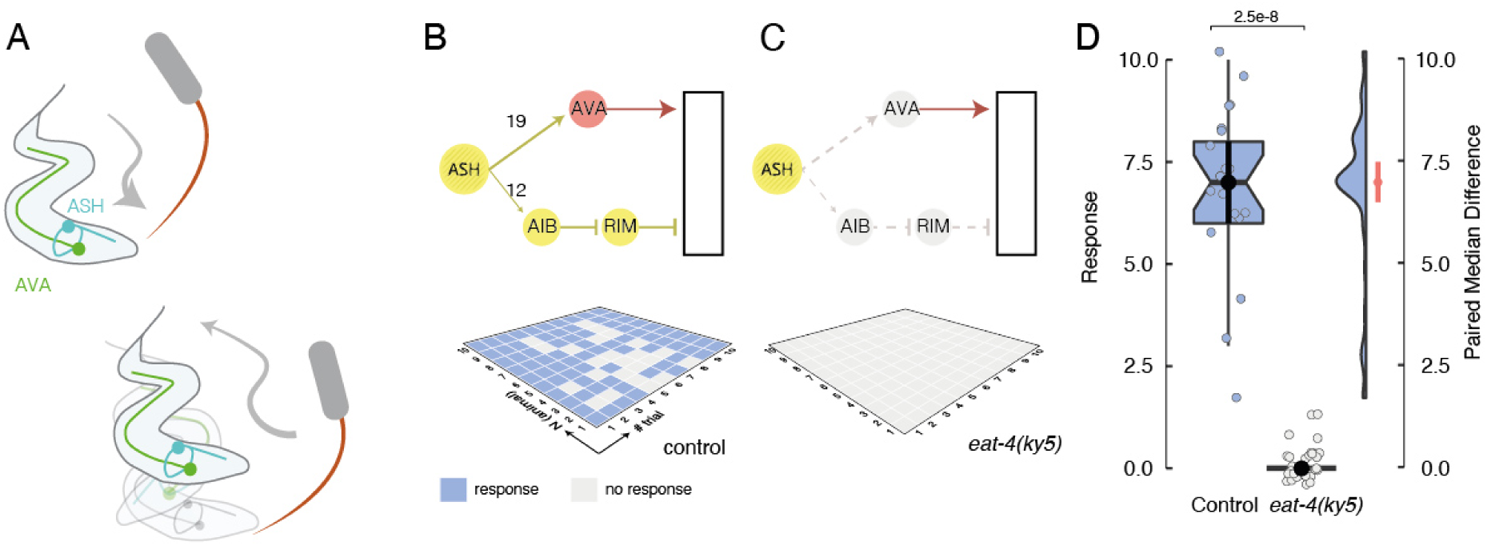
Nose touch mutants and transgenic. **A** Schematic of the behavioral assay. The yes/no response of a single animals is recorded as it navigates into a user-controlled obstacle (eyebrow hair). **B,C** Wiring of the nociceptive avoidance circuit with the representative result of an experiment of 10 trial conducted on 10 animals in (B) wildtype and (C) *eat-4(ky5)* mutants. Circuit nodes colored according to their neurotransmitter (yellow=glutamate; red=acteylcholine; grey=silent). Dashed arrow indicates broken connection in the *eat-4(ky5)* mutation. Grid plots show the yes/no response, color-coded according to its outcome, of individual animals as they navigated into the eyebrow hair. **D** Summary of results of both genotypes with a box plot encompassing 50% around the median and whiskers embracing 90% of the data. Floating axis on the right shows the bootstrapped distribution of their paired median difference (right axis), indicating the PMD *±* 95% confidence interval. *p*-value derived from a binary logistic regression, (see Methods).

**Figure S2.**
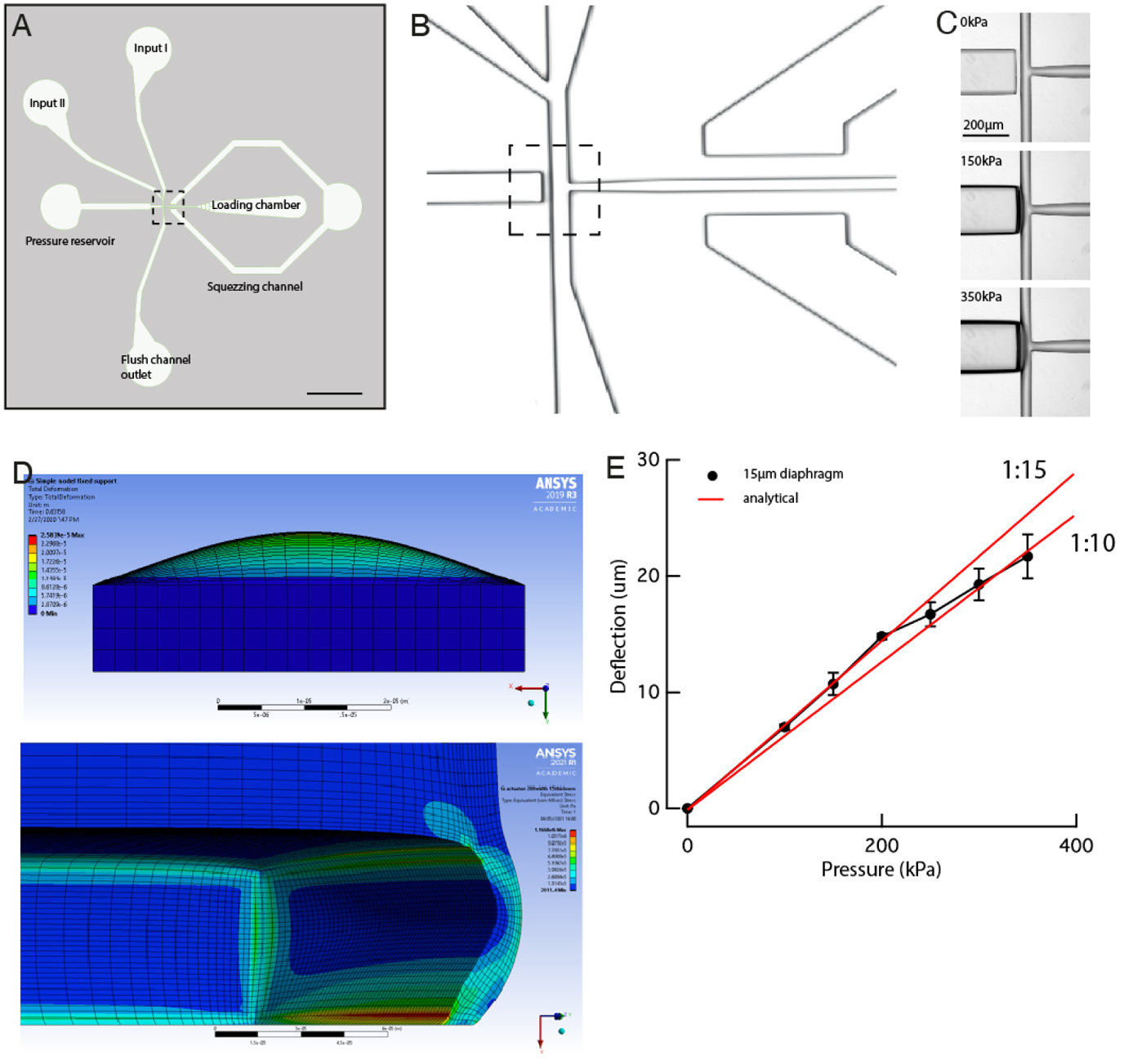
Design and calibration of the Trap and Slap design. **A** Layout of the microfluidic design. Scale bar = 1mm. **B** Photograph of the PDMS device of the dotted area shown in (A). **C** Photographs of the diaphragm deflection with increasing pressure on the channel of the dotted area shown in (B). **D** Finite element simulation of the plate deflection (top panel) showing the parasitic deformation of the bulk PDMS during the deflection. The bottom panel shows the stress contour in the device after inflation with xxx kPa back pressure. **E** Measurement of the 15*µ*m thick diaphragm deflection with increasing back pressure and the comparison to an analytical plate deflection model for two different PDMS mixing ratios 1:10 and 1:15 (red).

**Figure S3.**
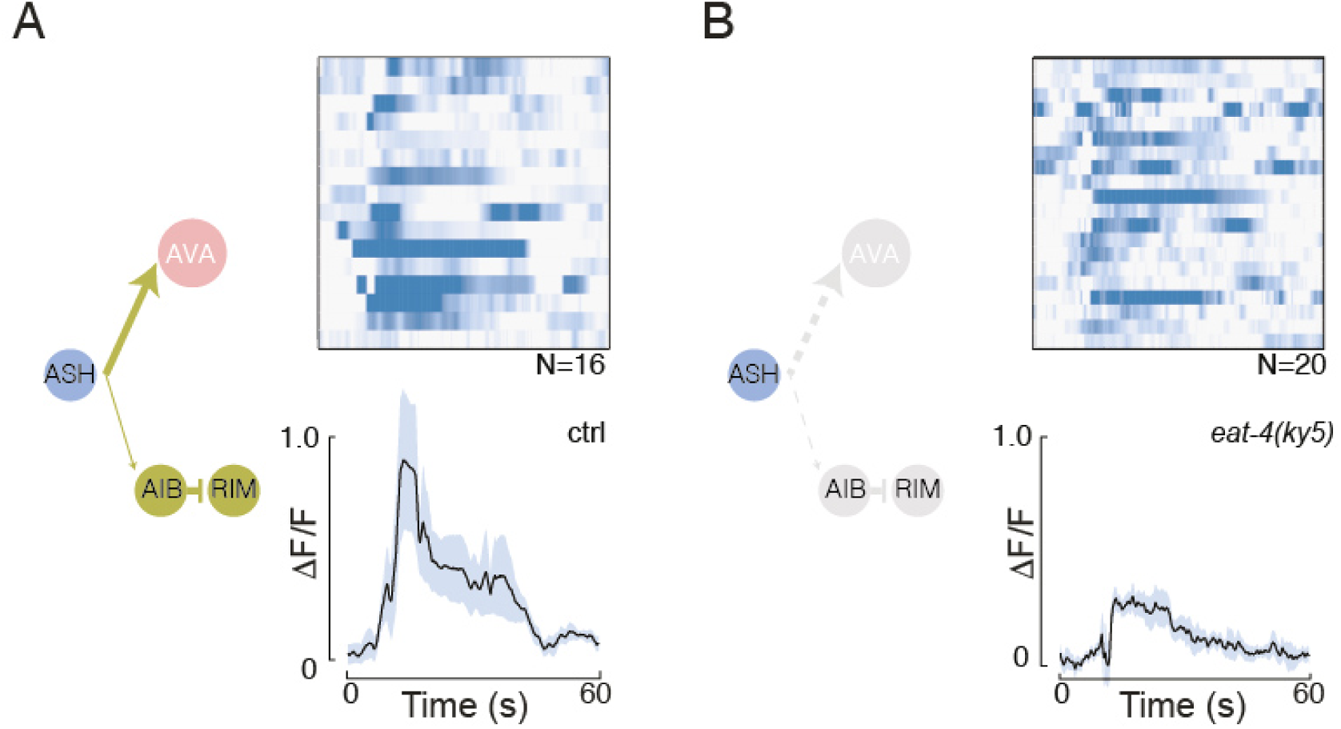
Mechanosensitivity of ASH does not depend on *eat-4*. **A,B** Schematic of the neuronal circuit with ASH highlighted in blue and glutamatergic edges in yellow and nodes colored corresponding their neurotransmitter usage (red, acteylcholine; yellow, glutamate). Stacked kymoraphs and average ASH:jRGECO1a fluorescence after pneumo-mechanical nose touch in the Trap’N’Slap device after 10s in (A) control and (B) *eat-4(ky5)* animals. Shaded area indicates SD around the mean (black trace). Broken connections in the *ky5* mutant are indicated as grey dotted lines.

**Figure S4.**
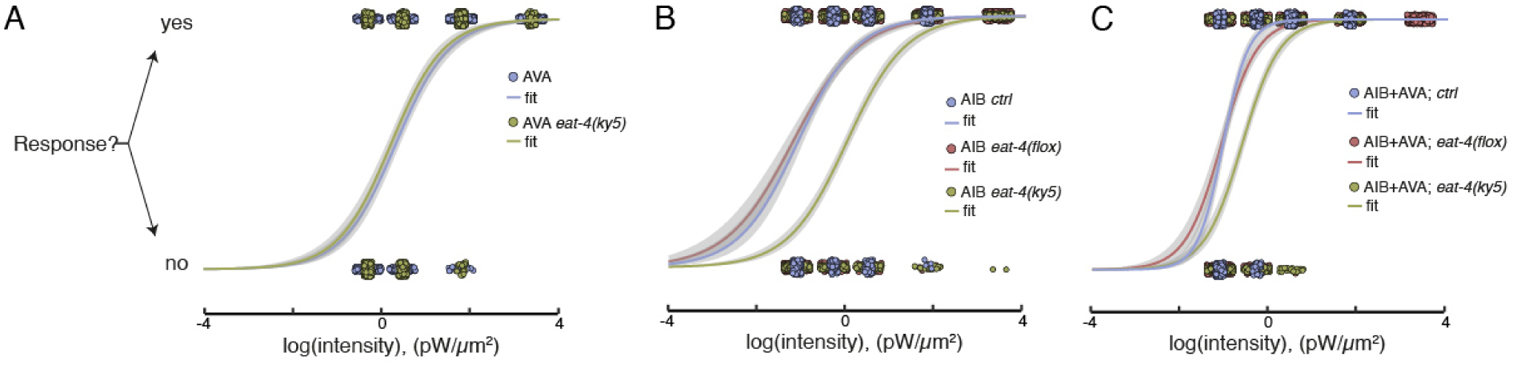
Efficiency of ChR2-HRDC to induce reversal behavior through AIB and AVA in the eat-4 neurotransmitter mutants. **A,B** Expression of ChR2-HRDC in (A) AVA and (B) AIB as a single copy transgene elicits behavioral responses independent of *eat-4*. **C** Combined expression of ChR2-HRDC in AVA and AIB elicits behavioral responses to low light and requires *eat-4* for full light sensitivity. Solid line corresponds to a binary logistic regression of the measured response as a function of the light intensity. N=30 animals, each tested 10 times.

**Figure S5.**
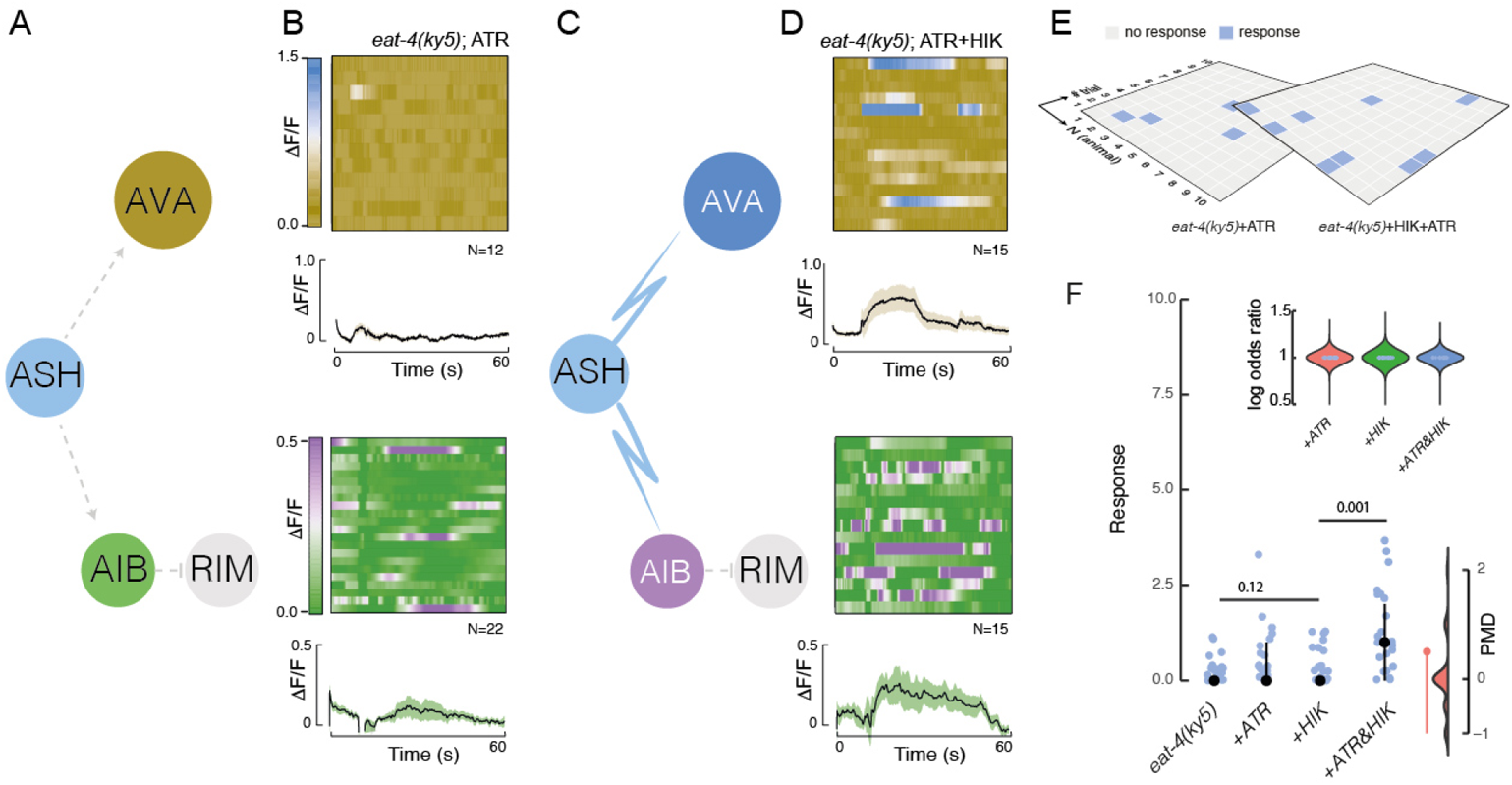
PhAST rescues calcium signaling but not behavior in the *eat-4(ky5)* mutation. **A,B** Schematic of the experimental circuit manipulation. Nodes are colored according to the LUT in (B), grey dotted edges indicate genetically perturbed synaptic connections in the *eat-4(ky5)* mutation. Bb) Image plot and average jRGECO1a fluorescence after pneumo-mechanical nose touch in the microfluidic device for AVA (khaki, blue) and AIB (olive, violet) in *eat-4(ky5)* animals expressing the light pathway in absence of the cofactors. Shaded area indicates SD around the mean (black trace). The vertical bar indicates the duration of the mechanical stress. **C,D** Schematic of the circuit with eNL expression in ASH (blue) and light-restored edges shown with blue arrows. (D) Image plot and average jRGECO1a fluorescence for AVA (khaki, blue) and AIB (olive, violet) for the same animals as in (B) but in presence of both cofactors. Shaded area indicates SD around the mean (black trace). **E** Behavioral response of *eat-4(ky5)* animals expressing the light pathway in presence and absence of all cofactors. **F** Summary of nose touch in *eat-4(ky5)* animals. Black dot and vertical bars indicate median *±* 95% confidence interval (CI). The p-value was derived from a Wald chi-squared statistics *z*^2^ = (*β*^^^*_j_*/SE(*β*^^^*_k_*))^2^, comparing the population measure of the response. The floating axis to the right indicates the bootstrapped distribution of the paired median difference (PMD) between HIKarazine single treated animals and double treated animals. Black point indicates median *±* 95%CI. Overlap of the CI with zero indicates low effect size and likely statistically insignificant distributions. Inset shows the log-odds ratio of finding a treated animal responding compared to the untreated mutant control.

**Figure S6.**
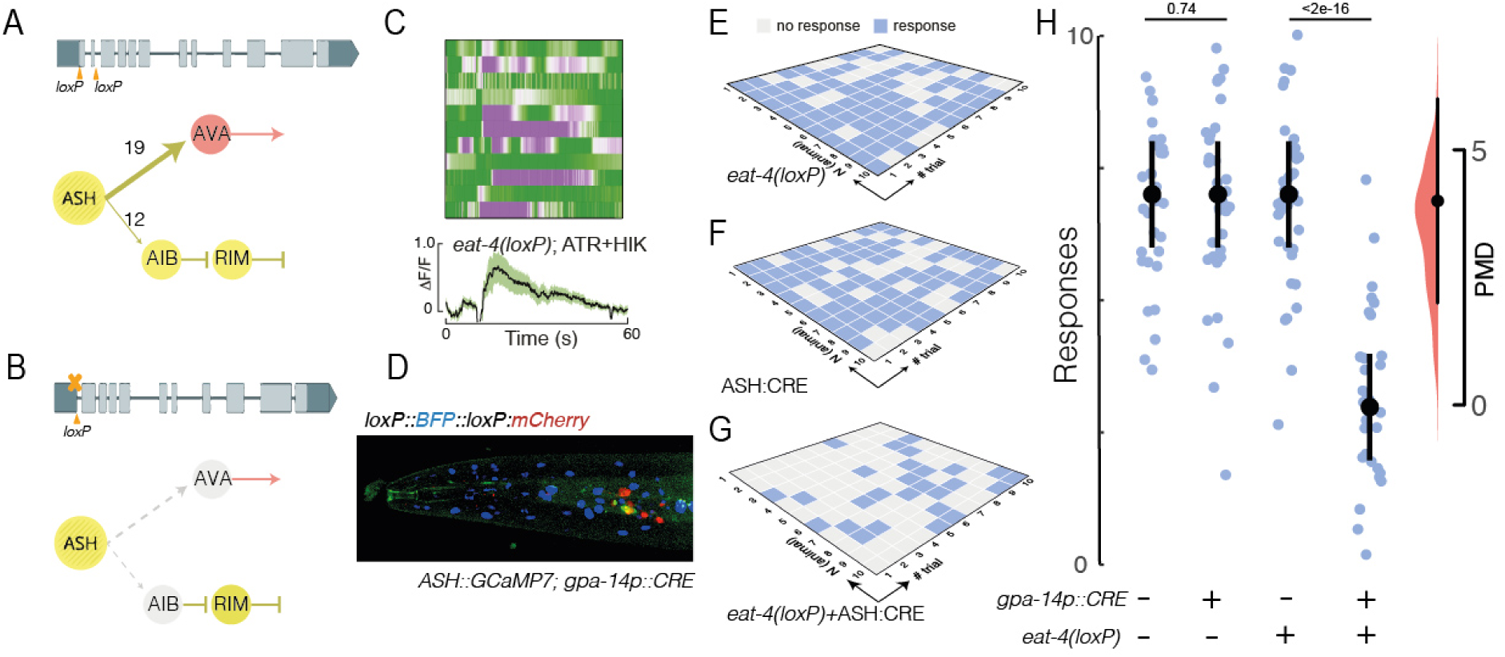
ASH-specific loss in glutamatergic signaling causes nose touch deficiency. **A,B** Schematic of the genetic strategy for ASH-specific ablation of *eat-4* and the location of the two lox sites flanking exons 1 and 2, (A) before and (B) after CRE recombination. The circuit indicates glutamatergic (yellow) and cholinergic (red) edges, with numbers of synaptic contacts above the arrow. **C** Stacked kymographs of individual RGECO videos and average AIB:jRGECO1a fluorescence after pneumo-mechanical nose touch in the microfluidic device in control and *eat-4(loxP)* animals in absence of CRE recombinase. Shaded area indicates SD around the mean (black trace). N=12 recordings. **D** Representative animal showing the split-CRE recombination pattern using a recombination reporter with a BFP-mCherry switch for successful recombinations coexpressing an sra-6p::GCaMP7 construct to highlight ASH (see also Fig. S9). **E-G** Grid plots showing a representative datasets of ten touches to ten animals for (E) *eat-4(loxP)* control, (F) *gpa-14p::CRE* single transgenics and (G) ASH(*eat-4*) loss-of-function animals after *gpa-14p::CRE* expression, color-coded according to its outcome (blue=positive response, grey=negative response to nose touch). **H** Summary of the nose touch response for the control and conditional *eat-4* knockout in ASH. Only for display purposes, a scatter of 10% was applied to each datapoint to avoid overlap. Circle indicates median, vertical bar indicates 95% confidence interval on the median. Floating axis indicates the paired median difference, derived from bootstrapping 100 independent distributions from the experimental data set. Black circle indicates median *±* 95%CI. Overlap of the CI with zero indicates low effect size and likely statistically insignificant distributions.

**Figure S7.**
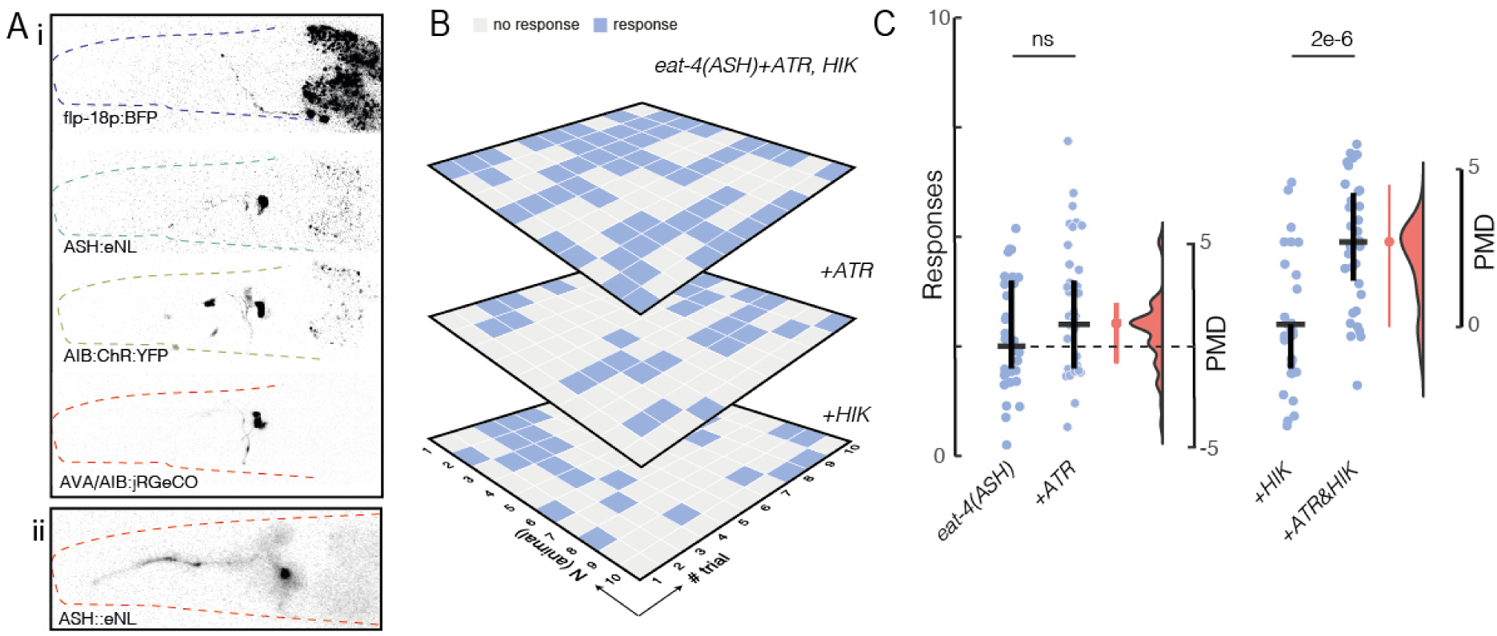
Nose touch response of soluble eNL. **A** i) Fluorescence micrograph of the individual transgenes used to express eNL in ASH and ChR2-HRDC in AIB and AVA. ii) Luminescence of the Nanolantern in ASH (compare to Figure 3). **B** Nose touch avoidance response of the conditional *eat-4*(loxP) mutant allele in ASH, coexpressing eNL in ASH and ChR2-HRDC in AIB and AVA supplemented with the cofactors indicated. **C** Summary of the scores for the nose touch experiment on 30 animals in all conditions tested. Only for display purposes, a scatter of 10% was applied to each datapoint to avoid overlap. Horizontal bar indicates median, vertical bar indicates 95% confidence interval on the median. Floating axis indicates the paired median difference, derived from bootstrapping 100 independent distributions from the experimental data set. Red point indicates median *±* 95%CI. Overlap of the CI with zero indicates low effect size and likely statistically insignificant distributions.

**Figure S8.**
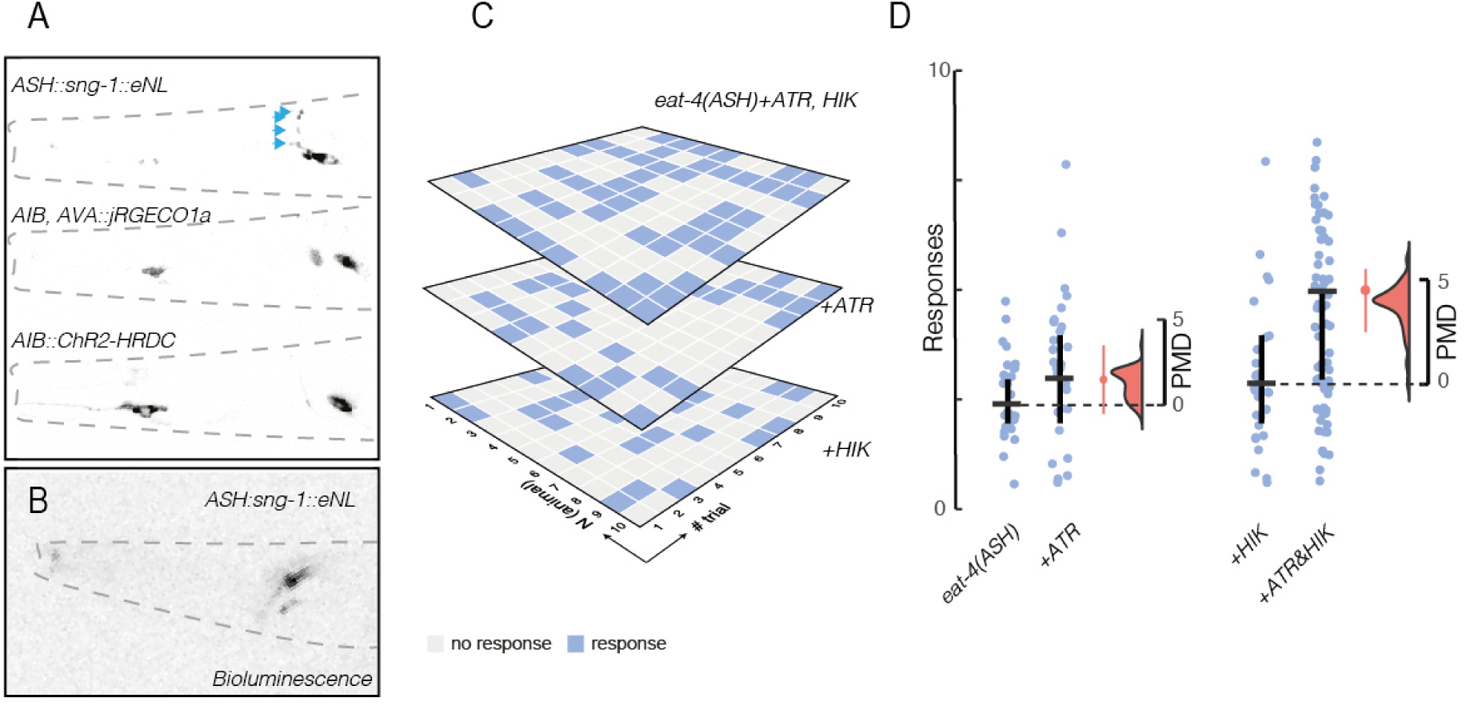
Nose touch response of synaptically localized eNL. **A** Fluorescence micrograph of the synaptically localized Nanolantern, fused to sng-1 synaptogyrin and the other transgenes. Cyan arrowheads point towards the presumptive synapses indicated by high sng-1 intensities. **B** Calcium saturated bioluminescence micrograph of the sng-1::eNL **C** Nose touch avoidance response of an animal carrying the ASH specific *eat-4(loxP)* mutant allele in absence and presence of indicated cofactors, coexpressing synaptic *sng-1*::eNL in ASH and ChR2-HRDC in AIB and AVA, displayed as a grid plot, colorcoded according to its outcome (blue=positive response, grey=negative response to nose touch). **D** Summary of the scores for the nose touch experiment on 30 animals in all conditions tested. Only for display purposes, a scatter of 10% was applied to each datapoint to avoid overlap. Horizontal bar indicates median, vertical bar indicates 95% confidence interval on the median. Floating axis indicates the paired median difference, derived from bootstrapping 100 independent distributions from the experimental data set. Red point indicates median *±* 95%CI. Overlap of the CI with zero indicates low effect size and likely statistically insignificant distributions.

**Figure S9.**
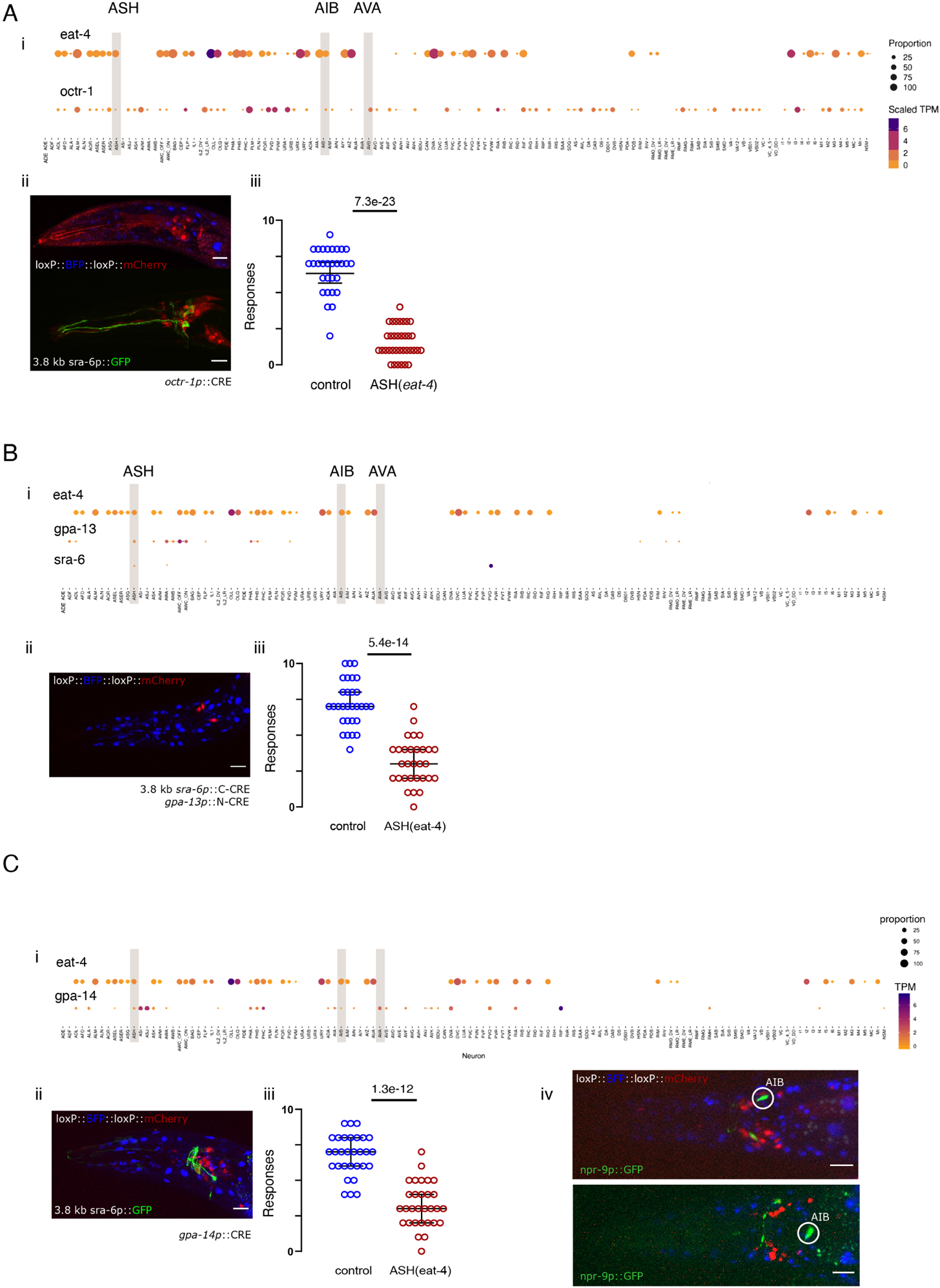
CRE expression under the control of differe5n3t promoters. **A** Expression of CRE under the control of the *octr-1* promoter. (i) Expression pattern of eat-4 and octr-1 as determined in Ref (*37*) with overlap highlighted in ASH, AIB and AVA. (ii) CRE-loxP recombination was tested in a color switch strain that expresses nuclear BFP in absence of CRE activity and nuclear mCherry in the cells where CRE is active. Recombination was visible in ASH and two other cells as judged by coexpression with a sra-6p:GCaMP transgene known to drive in ASH (*23*). (iii) Outcome of nose touch assays in worms with the floxed *eat-4* allele and expression of *octr-1p*::CRE. **B** Expression of a split CRE to establish ASH-specific expression (see Methods). (i) Expression pattern of *eat-4*, *gpa-13* and *sra-6* as determined in Ref. (*37*) with overlap highlighted in ASH, AIB and AVA. (ii) CRE-loxP recombination pattern showing successful BFP*>*mCherry switch in cells in which the two promotors intersect and thus CRE is activity is reconstituted. (iii) Nose touch response of animals with a floxed *eat-4* allele and expression of the split *sra-6p*::C-CRE and *gpa-13p*::N-CRE. **C** Expression of CRE under the control of the *gpa-14* promoter. (i) Expression pattern of *eat-4* and *gpa-14* as determined in Ref (*37*) with overlap highlighted in ASH, AIB and AVA. (ii) CRE-loxP recombination was tested in a color switch strain that expresses nuclear BFP in absence of CRE activity and nuclear mCherry in the cells where CRE is active. In addition, coexpression of mCherry with the ASH specific 3.8 kb *sra-6p* driving GFP expression was tested. (iii) Outcome of nose touch assays in worms with the floxed *eat-4* allele and expression of *gpa-14p*::CRE. Scale bar= 15 *µ*m. p-value corresponding to an unpaired, parametric t-test with 95% confidence interval. Median and 95% confidende interval are depicted in all dot plots. (iv) Representative images of a CRE-activity reporter animal expressing gpa-14p::CRE and npr-9p::GFP to highlight potential recombination in AIB. As can be seen by the absence of the GFP/mCherry overlap, gpa-14p::CRE does not drive recombination in AIB. Two different animals are representative for 8 randomly picked animals.

## 6 Supplementary Videos

**Video S1: Nociceptive avoidance behavior.** Representative video of an animal navigating into an obstacle in (A) wildtype and (B) *eat-4(ky5)* background.

**Video S2: Pneumatic stimulation of a trapped animal inside the Trap’N’slap device.** Representative video of a wildtype animal subjected to a 2.5bar stimulus, recorded in brightfield microscopy.

**Video S3: Calcium imaging under mechanical stimulation in ASH** Representative video of the fluorescence intensity of (A) control and (B) *eat-4(ky5)* mutant animals expressing jRGECO1a in ASH. Animals were immobilized in the microfluidic device during the presentation of a 2s mechanical stimulus after 10s. Scalebar=30*µ*m, framerate=10Hz. Anterior to the left. Same color LUT as in figure 1.

**Video S4: Calcium imaging under mechanical stimulation in AVA** Representative video of the fluorescence intensity of (A) control and (B) *eat-4(ky5)* mutant animals expressing jRGECO1a in AVA. Animals were immobilized in the microfluidic device during the presentation of a 2s mechanical stimulus after 10s. Framerate=10Hz. Anterior to the left.

**Video S5: Calcium imaging under mechanical stimulation in AIB** Representative video of the fluorescence intensity of (A) control and (B) *eat-4(ky5)* mutant animals expressing jRGECO1a in AIB. Animals were immobilized in the microfluidic device during the presentation of a 2s mechanical stimulus after 10s. Scalebar=40*µ*m, framerate=10Hz. Anterior to the left.

**Video S6: Optogenetic stimulation of AVA** Representative video of a reversal response to optogenetic stimulation of an animal expressing ChR2-HRDC in AVA in presence (right) and absence (left) of the photosensitizer all-trans retinal (ATR).

**Video S7: Optogenetic stimulation of AIB** Representative video of a reversal response to opto-genetic stimulation of an animal expressing ChR2-HRDC in AIB in presence (right) and absence (left) of the photosensitizer ATR.

**Video S8: Crawling animal with calcium sensitive Nanolantern reporting Body wall muscle activity** Representative video of a freely crawling animal expressing a calcium sensitive Nanolantern in body wall muscles. Increases in intensity on the concave side of the body bend indicates that the calcium influx increases quantum yield of the Nanolantern probe.

## 7 Tables

**Table S1: Avoidance behavior to nose touch** Summary of the outcome to nose touch of the different strains used in this study.

**Table S2: AVA and AIB Response to blue light** Raw data of the optogenetic experiments conducted in the several strains used.

**Table S3: Strains** Summary and characteristics of strains appearing in figures (sheet 1) and other strains used in this study (sheet 2).

**Table S4: Plasmid name and DNA sequences** Plasmids (sheet 1) and DNA sequences (sheet 2) used in this study.

**Table S5: CRISPR and mutagenesis sequences** Compilation of the crRNAs, homology repair templates used in CRISPR/Cas9 edits and primers used for directed mutagenesis.

